# Integrative Structure and Function of the *Vibrio cholerae* Competence Pilus Machine

**DOI:** 10.1101/2025.10.14.680967

**Authors:** Stefano Maggi, Stefan Kreida, Lixinhao Yang, Abigail E. Teipen, Diane L. Lynch, Ankur B. Dalia, James C. Gumbart, Grant J. Jensen

## Abstract

Type IV pili are hair-like, surface-exposed polymers involved in many fundamental biological processes. The competence pilus (CP) is responsible for natural transformation, the ability to acquire exogenous DNA. Here we purified the *Vibrio cholerae* CP and used electron cryo-microscopy (cryo-EM) to obtain a 3.3 Å map of the fiber by helical reconstruction. We then used electron cryo-tomography (cryo-ET) to reveal the *in-situ* architecture of the competence pilus machine (CPM). Compared to other type IVa pilus machines, the CPM has unique characteristics including multiple conformational states, a ring-like density formed by PilQ’s C-terminal domain absent in the previously solved SPA structure, and the presence of a double ATPase ring in the cytoplasm. Using integrative modeling with the cryo-ET map as an envelope, we generated a full-length pseudoatomic model of the CPM. Molecular dynamics (MD) simulations of the entire CPM embedded within realistic bacterial membranes and the peptidoglycan cell wall revealed that the model demonstrates structural integrity under physiological conditions. Steered MD simulations of pilus extension and retraction through PilQ revealed gate-opening mechanisms and demonstrated the channel’s mechanical resistance and flexibility during translocation.

## Introduction

Horizontal gene transfer is one of the most powerful driving forces of bacterial evolution, and type IVa pili (T4aP) are implicated in all three classic gene transfer mechanisms: transformation^1^, transduction^2^, and conjugation^3,4^. One of the best-characterized T4aP systems is the *Vibrio cholerae* competence pilus (CP), a multi-subunit megadalton machine required for natural transformation^5^. Competence requires a minimal set of 19 genes that can be grouped into three functional classes: proteins that form the CP fiber, proteins that assemble the competence pilus machine (CPM), and proteins needed for DNA recombination^5^.

Structurally, type IVa pili are composed of two parts: the tip and the pilus filament^6^. The tip is directly involved in DNA binding^7,8^ and is believed to be made of a collection of minor pilin proteins^6,9,10^. Though the structures of individual minor pilins have been solved^6^, a complete atomic model of the tip is still missing^5^. In contrast, several T4aP filament structures have been solved^11–14^. These structures usually consist of multiple copies of a single protein (the major pilin) arranged in a helical pattern. Major pilins typically have a “lollipop” fold^15^ formed by a C-terminal globular domain packed against an N-terminal α-helix of approximately 50 residues, and are characterized by the presence of an unusual disordered central region contributing to the intrinsic flexibility of the pilus filament^16^. The CP major pilin is PilA, and its assembly into a filament (extension) and disassembly (retraction) are mediated by the CPM.

In *V. cholerae*, 10 proteins make up the CPM: PilB, PilC, PilM, PilN, PilO, PilP, PilQ, PilT, PilU, and VC1612. PilB and PilT can each antagonistically bind the machine and allow for extension (PilB) or retraction (PilT) of the pilus^17^. PilU is important for forceful retraction^17^, but its activity requires PilT^17,18^. In the absence of PilT and PilU, the CP can slowly retract via a motor-independent mechanism of spontaneous depolymerization of the PilA monomer. This depolymerization is likely due to the intrinsic instability of the pilus fiber^19,20^. A previous study from our laboratory reported the structure of PilQ at high resolution^21^. PilQ is a secretin, which forms the outer membrane (OM) pore and acts as a gate for the pilus fiber while ensuring OM integrity. Secretins of type IV pili are often associated with TsaP (T4P secretin-associated protein), a bimodular protein that forms a ring around the secretin and anchors it to the peptidoglycan (PG) layer using TsaP’s N-terminal LysM domain^22^. In *Vibrio cholerae*, the gene VC0047 encodes a TsaP homolog, the absence of which reduces transformation efficiency ∼10-fold^23^.

The functions of the other CPM genes have been inferred by homology^24^: PilN, PilO, and PilP may work together as inner-membrane (IM) bound periplasmic adaptors that align the secretin with the machine’s cytoplasmatic components. PilC and PilM could be the IM rotor and the cytoplasmic stator, respectively. VC1612 might act as a pilotin by helping facilitate PilQ secretin assembly^25^.

To gain insight into how all these components assemble and interact, we purified the CP fiber, imaged it with electron cryo-microscopy (cryo-EM), and solved its structure by helical reconstruction to near-atomic resolution. We also investigated the CPM architecture through electron cryo-tomography (cryo-ET) of whole cells, subtomogram averaging, and unsupervised classification. Our data revealed different CPM conformations in piliated and non-piliated machines. We also observed that the *in-situ* PilQ AMIN domain forms a ring-like structure absent in the previously solved *Vc*PilQ single particle map^21^. We further discovered that PilT and PilU form a double ATPase ring at the base of the machine. Using this data, integrative modeling, and AI protein structure prediction tools, we built a pseudoatomic integrative model of the entire CPM. Based on this model, we performed molecular dynamics (MD) simulations to elucidate molecular rearrangements during pilus extension and retraction.

## Results

### PilA adopts a canonical bacterial T4aP fold

*Vibrio cholerae* encodes three distinct type IVa pilus machines: the toxin-coregulated pilus, the mannose-sensitive hemagglutinin pilus, and the CP. To investigate the structure of the CP and the architecture of the CPM, we used a *V. cholerae* strain lacking both toxin-coregulated and mannose-sensitive hemagglutinin pilus systems. We placed TfoX, *V. cholerae*’s main activator of natural competence, under the control of an IPTG-inducible promoter and knocked out *pilT*, which resulted in a hyper-piliated aggregating cell phenotype^21–23^. We purified the pili through a series of low-speed centrifugations at pH 9.5, taking advantage of the fact that these pili form dense bundles at pHs below 10 and dissociate into single filaments at pHs above 10. Purified pili were exchanged into 150-mM ethanolamine, pH 10.6, for vitrification. Cryo-EM movies were collected on a 300 kV Krios microscope with a final pixel size of 0.832 Å. Helical reconstruction was done in Relion^27^. Start- and end-coordinates of individual filaments were defined in motion-corrected micrographs (Fig. 1A), from which we extracted 77,000 300x300-px boxes with a 10 Å overlap, which roughly corresponds to the helical rise of the filament. To avoid bias, initial helical parameters were obtained by running a round of 3D classification with a cylinder 80 Å in diameter as a starting reference without applying helical symmetry. The estimated twist and rise were extracted from the best resulting class and used as starting parameters for subsequent 3D classifications and refinements with C1 symmetry (one-start helix). The final reconstruction was obtained from 22,000 boxes and had a final resolution of 3.3 Å (map:map FSC at 0.143; Fig. S1), a rise of 10.75 Å, a twist of 94.65°, and a width of 67 Å (Fig. 1). The helical parameters fall within the range of other T4aP family members for which structural information exists^28^.

**Figure 1.**
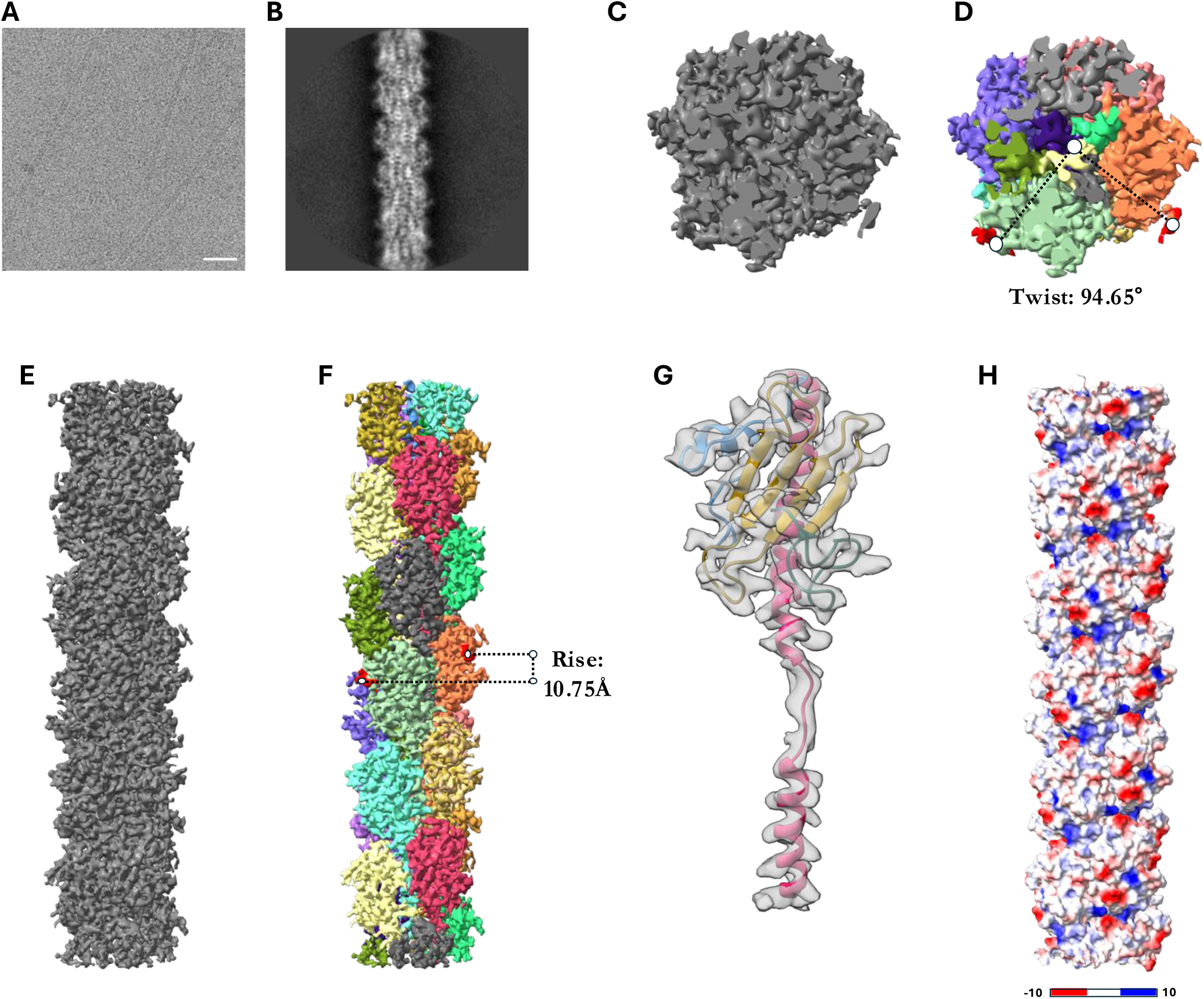
Helical reconstruction of the competence pilus. A) Raw micrograph of frozen hydrated pili. B) 2D class example used for *de novo* 3D model generation. C) Top view of the CP map after the final 3D refinement colored by chain identity (D) highlining the twist of 94.65 Å. E–F) Side view of the same map as C–D highlighting the 10.75 Å rise. G) Extracted map of a CP protofilament with the atomic model (pink : N-terminal alpha helix with melted region, blue: alpha-beta loop, yellow: beta-sheet, gray: D region). H) CP atomic model colored by Coulombic potential (kcal·mol^-1^·e^-1^ at 298°K) showing positive and negative patches along the helical symmetry.

Like other T4aP pilins, we found the *V. cholera* PilA structure to be characterized by a “lollipop-like” fold with a surface-exposed C-terminal globular domain, mainly composed of a beta sheet, folded against an alpha-helical N-terminal domain (Fig. 1G). In pili, the N-terminal domain is buried inside the CP filament (Fig. S2A) and contains a molten region evident in the map density (Fig. 1G). The N-termini are positioned at the core of the structure. Extra side-chain density was observed for Phe 1, which could be assigned to an N-terminal methylation site (Fig. S2B)^29^. The relative orientation in the final model (methyl group up, phenyl group down) was based on packing restrictions, as the opposite orientation resulted in severe steric clashes during refinement in Phenix. A small, solvent-exposed helix (residues 77–81) links the N-terminus with the C-terminal globular domain. The C-terminal domain is composed of four antiparallel beta strands that orient hydrophobic residues toward the pilus core and polar residues toward the solvent (Fig. S2C). It also includes a C-terminal loop (residues 136–153) in front of the beta sheet. This orientation is further stabilized by a disulfide bond between conserved Cys 133 and Cys 146 on the C-terminal loop. The C-terminal Asn 153 could be stabilized by the nearby presence of Gln 122 and several positively charged residues (Lys 108 and 120), although this is not evident from the density map (Fig. S2D).

### The pilus filament is characterized by charged patches

CP bundling can facilitate kin recognition in *V. cholerae*. We found that the pilus filament contains many charged, surface-exposed residues in electropositive and electronegative patches along its helical symmetry (Fig. 1H). We speculated this could explain the propensity of hyperpiliated Δ*pilT* cells to aggregate. Because the pili bundling phenotype switches around a pH of 10, we thought it likely that one or more lysine residues could be involved. We therefore investigated the role of three surface-exposed lysines on this aggregation phenotype by mutating them to serine (*pilA*_K40S_, *pilA*_K108S_, *pilA*_K120S_) (Fig. S3). Strains expressing *pilA*_K40S_ and *pilA*_K108S_ did not aggregate in solution (Fig. S3B) but retained WT levels of transformation, indicating that functional CP were still generated in this background. We were also able to visualize pili after staining cells containing the cysteine knock-in *pilA*_S67C_ with 488-malemide (Fig. S3C), which further supports that these mutations do not disrupt CP assembly. We conclude that PilA^K40^ and PilA^K108^ are involved in inter-pili interactions.

### Subtomogram averaging of the CPM reveals distinct structural states

In *V. cholerae* tomograms we saw classical components of bacterial cells like storage granules, flagellar motors, type VI secretion system tubes, and, given the hyper-piliated phenotype, several CP bundles (Fig. 2A). Piliated and non-piliated machines were easily identifiable in the bacterial envelope by their secretin densities in the outer membrane and their ring densities in the periplasm. To increase the signal-to-noise ratio (SNR), we averaged 349 non-piliated and 604 piliated machines in PEET^30^. Initial subtomogram averages (STAs) showed a characteristic T4aP architecture with three main densities: the OM pore, mid-periplasmic ring, and lower periplasmic ring. In addition to these densities, the CPM contains a novel fourth ring encircling the OM pore (Fig. 2B and C), which we have called the flanking ring. In both piliated and nonpiliated STAs, the lower periplasmic ring is less defined compared to the other densities, which could be the result of structural heterogeneity (Fig. 2B and C). We therefore classified the particles using principal component analysis (PCA) followed by K-means clustering^31^.

**Figure 2.**
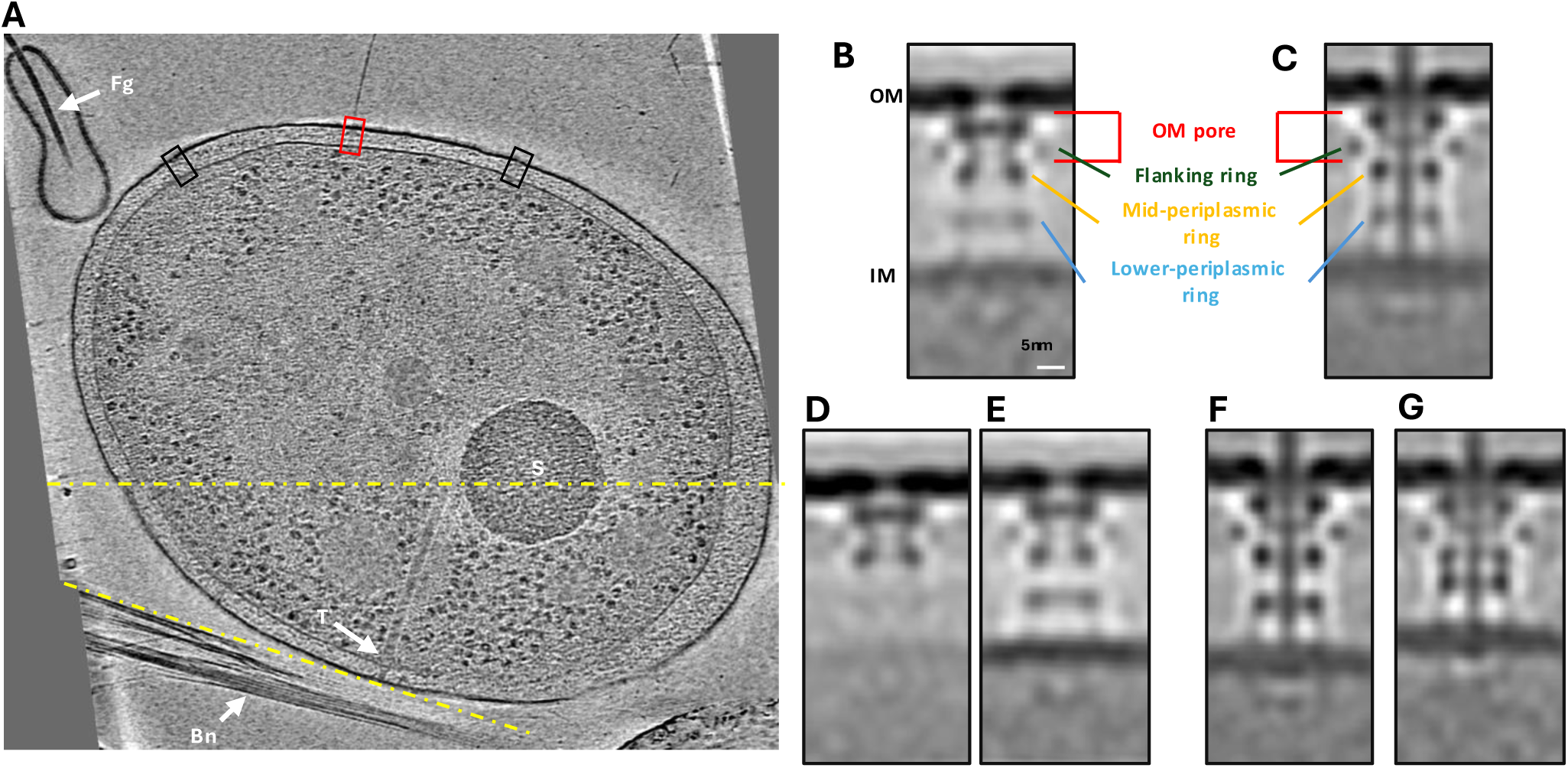
Cryo-ET of the ΔPilT strain reveals different CPM states. A) Composite image of different planes of a representative *Vibrio cholerae* tomogram showing the flagellum (Fg), competence pilus bundles (Bn), a type VI SS (T), a storage granule (S), and type IV piliated (red box) and non-piliated (black boxes) machines. B–C) Subtomogram averages of the non-piliated (349 particles) and piliated (604 particles) CPMs showing a faint lower periplasmic ring. D–E) Classification of the non-piliated machines results in two classes at different assembly states of the lower periplasmic ring. F–G) Classification of the piliated machines results in two classes showing different conformational states of the lower periplasmic ring.

In the non-piliated dataset, we identified two distinct classes. One class showed only the OM pore and its flanking densities (231 particles, Fig. 2D), while the other also exhibited a well-defined lower periplasmic ring (210 particles, Fig. 2E). The T4aP machine assembles from the outside in^32^, with the secretin assembling in the OM prior to the formation of the lower periplasmic ring. Thus, we hypothesize these two classes could represent machines in different assembly stages.

Classification of the piliated state also revealed two distinct classes. The major class (509 particles) had a distance between the IM and OM of 37 nm (Fig. 2F), and the minor class (95 particles) showed a reduction in space between the IM and OM (32 nm) with the mid- and lower periplasmic rings in close contact (Fig. 2G). To ensure these were distinct conformational states of the CPM rather than a reflection of a distinct bacterial sub-population (e.g., persister cells) in our sample, we made sure these two conformations were both found in individual cells. Because 100 of 292 cells had both architectures, we conclude that the compacted state is a conformational state and does not reflect a low-frequency phenotype (Fig. S4). Additionally, neither class contained a clear PilB cytoplasmic density at the base of the pilus machine like those that have been seen in the *M. xanthus* T4aP^33^ and the *V. cholerae* toxin-coregulated pilus^34^ systems. This absence suggests a low PilB occupancy^5^ in the CPM.

### The secretin PilQ is tethered to the peptidoglycan by AMIN domains

STAs of both piliated and non-piliated machines displayed a novel ring flanking the OM pore density. PilQ homologs of previously characterized T4aP systems either don’t have an AMIN domain (*T. thermophilus*, *V. cholerae* toxin-coregulated pilus) or possess three AMIN copies (*M. xanthus*). Uniquely, *Vc*PilQ has only one AMIN domain separated from the beta-barrel pore via flexible linkers^21^ that anchor the secretin to the peptidoglycan layer^35^. Thus, we hypothesized that the novel flanking ring could be formed by this AMIN domain. To test our hypothesis, a GFP tag was introduced at residue 35 of PilQ, just before the AMIN domain (residues 44–147). The STA of the piliated PilQ_35_sfGFP machine showed an additional density corresponding to the size of sfGFP next to the flanking ring, confirming its AMIN identity (Fig. 3, Top).

**Figure 3.**
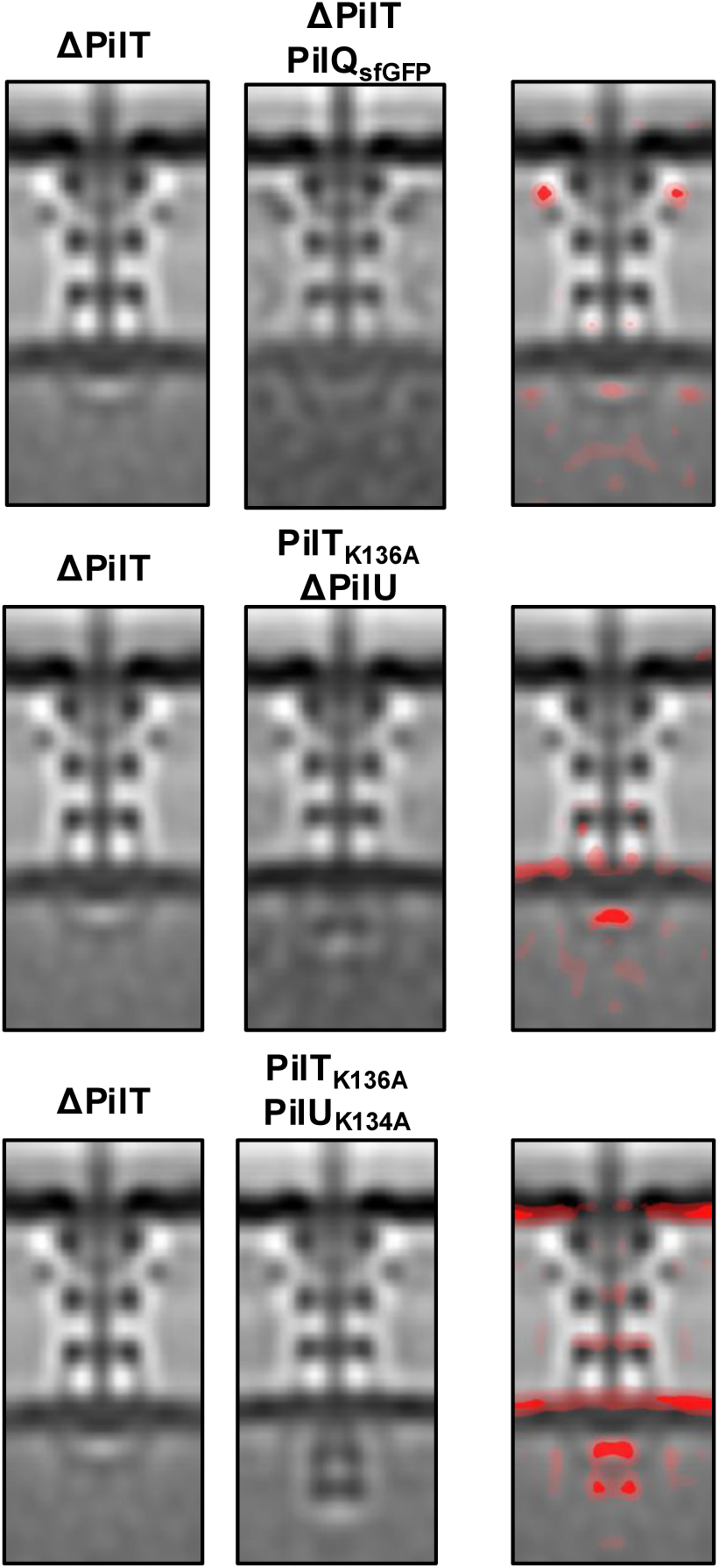
Localization of PilQ AMIN domain, PilT, and PilU in the piliated STA. The piliated ΔPilT STA (left column) was used as a reference for PilQ, PilT, and PilU mutants (center column) to calculate difference maps (right column). Top) STA of the ΔPilT PilQ_sfGFP_ strain revealing the position of the AMIN domain. Middle) STA of the PilT_K136A_ ΔPilU strain showing the location of PilT at the base of the machine. Bottom) STA of the PilT_K136A_ PilU_K134A_ strain revealing the PilT-PilU complex.

### PilT localizes to PilU

We next investigated the localization of PilT. Given the almost complete absence of pili on cells with PilT or PilU^17^, we collected tomograms of strains with a point mutation in the ATP binding site (PilT_K136A_) in the Δ*pilU* background, thus preventing pilus retraction. The STA of the *pilT*_K136A_ Δ*pilU* strain revealed an additional small density at the base of the CPM, confirming previous evidence for PilT localization at the base of the machine^33^ (Fig. 3, Middle). Given the dependence of PilU on PilT, we investigated PilU localization by imaging an inactive form (*pilU*_K134A_) in the *pilT*_K136A_ background. In the piliated STA of this strain, we saw a clear additional density just beneath the PilT density, revealing that PilT and PilU stack in the cytoplasm (Fig. 3, Bottom).

### Competence pilus machine model

To construct a complete model of the CPM (Fig. 4), we annotated each gene involved in piliation with its predicted localization, domain organization, and the availability of homologous solved structures (Fig. S5; for a full description, see supplementary material). In parallel, we built full-length monomeric models of each protein by using AlphaFold^36^ and ColabFold-multimer^37^ to predict protein complexes. In addition to a careful inspection of model statistics, we checked prediction quality by directly superimposing these models with solved structures of homologs (*Neisseria meningitis* PilQ-PilP, PDB: 4AV2; *Pseudomonas aeruginosa* PilN-PilM, PDB: 5EOU; and *V. cholerae* EspL-EspE complex for PilM-PilB, PDB: 2BH1).

**Figure 4.**
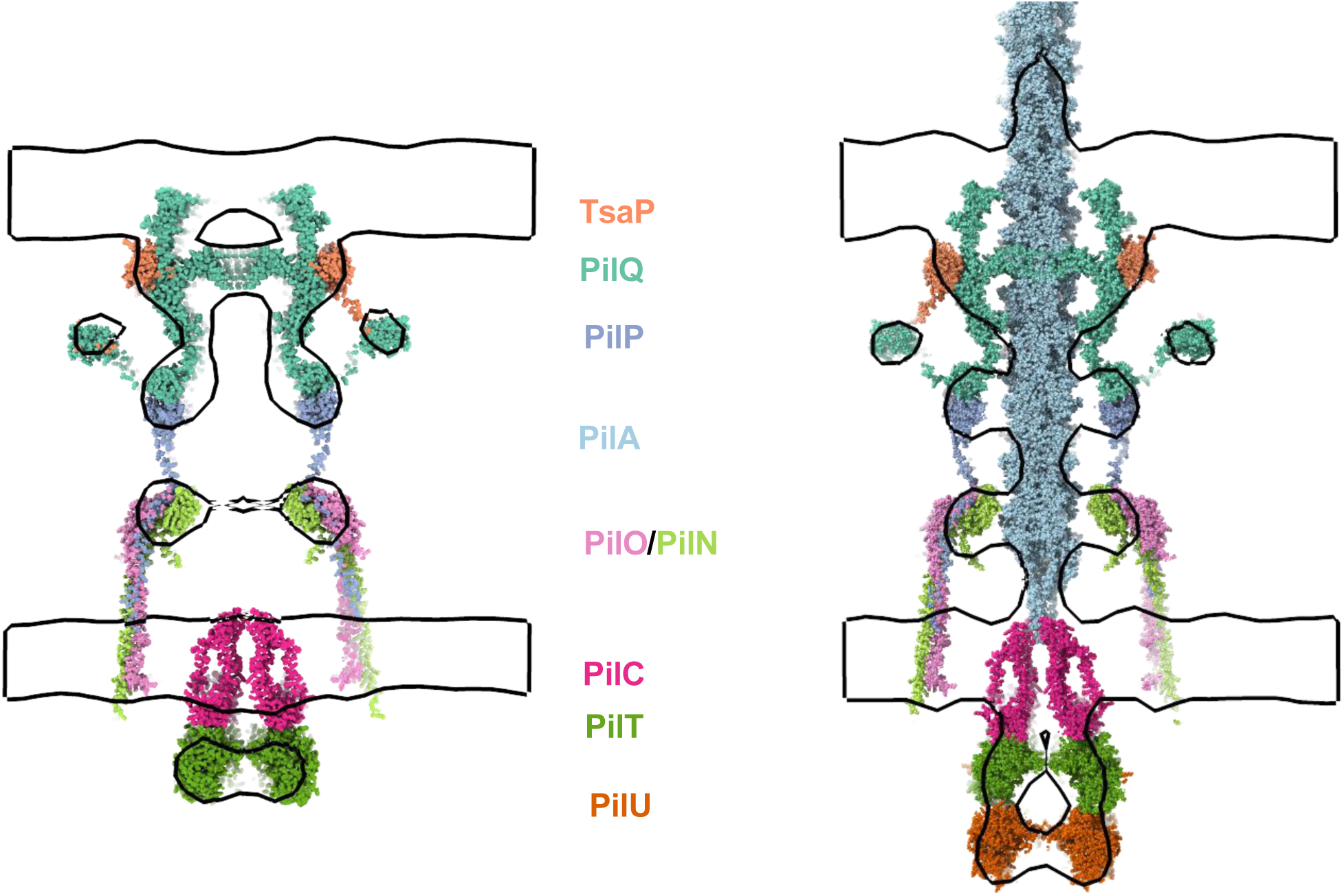
Integrative model of the CPM. Left: central slice. Right: side view. STA envelopes are shown in black outline.

PilQ is the only *V. cholerae* CPM protein with a solved structure; therefore, we used it as our starting point for modeling. To model the PilQ-TsaP complex we used molecular docking with the *Pseudomonas aeruginosa* PilQ-TsaP complex as a template (PDB: 6VE2, Fig. S6–7). When modeling the PilQ-PilP complex, the best result was obtained using *Nm*PilQ-PilP as a template followed by docking in HADDOCK^38,39^ using conserved residues at the protein-protein interface as interacting partners (Fig. S8–10). For the PilN-O-P prediction, its ColabFold prediction matches well with the experimentally characterized *Pa*PilN-PilO coiled coil^40,41^ (Fig. S11–12). Similarly, our ColabFold PilN-PilM (Fig. S13) and PilM-PilB (Fig. S14, 15) models are consistent with the solved structures of *Pa*PilN_1-12_-PilM and *Vc*EspL-EspE, respectively.

PilN is directly involved in PilM recruitment in the cytosol. In our model, PilN faces inward near the pilus fiber in the periplasm. Given its organization, PilN could be a sensor able to “feel” the pilus in the machine and transmit the signal to PilM.

Based on the current mechanistic model of T4aP motors, PilB and PilT induce different rotations of PilC (clockwise and counterclockwise, respectively)^42^. We modeled all three ATPases (PilB, PilT, and PilU) as hexamers in accordance with existing literature^17,42,43^. The type IV IM platform PilC is generally assumed to be a dimer^42,44–46^, so we modeled it accordingly, but it should be noted that the *Klebsiella pneumoniae* PulF protein (a homolog of PilC) in a type II secretion system has recently been shown to form trimers^47^; therefore, it is not clear if members of the PilC family form dimers, trimers, or both. The PilC_2_PilB_6_ and PilC_2_PilT_6_ models had similar confidence (ipTM) scores (Fig. S16, S18), but only PilT has been extensively characterized. The PilT AIRNLIRE motif is known to be important for pili retraction but is not involved in ATP hydrolysis^48^, and in our model, the motif is at the PilT-PilC interface (Fig. S17). Because this interface is not involved in ATP hydrolysis, the motif’s placement supports the validity of the model. PilB’s orientation in our PilC_2_PilB_6_ model has the second N-terminal domain facing PilC, which is similar to the previously proposed model^42^. To gain more insight on how the homologous proteins PilB and PilT can lead to different rotations, we superimposed the models on the ATPase domain and found that the homologs likely engage PilC with two opposite orientations (Fig. S19): PilB’s N-terminal domain binds PilC to trigger extension while PilT’s C-terminal domain binds PilC to trigger retraction.

### CPM structure after relaxation

To explore the structural dynamics and stability of the CPM model, we performed MD simulations. We constructed an all-atom model to mimic the CPM in its native environment, incorporating the entire assembly, realistic outer and inner membranes, and the PG cell wall, totaling approximately 4.9 million atoms (Fig. 5A). A 120-ns MD simulation revealed that the CPM model maintained large-scale structural stability with no observable breakdown, suggesting the proposed assembly is robust under physiological conditions.

**Figure 5.**
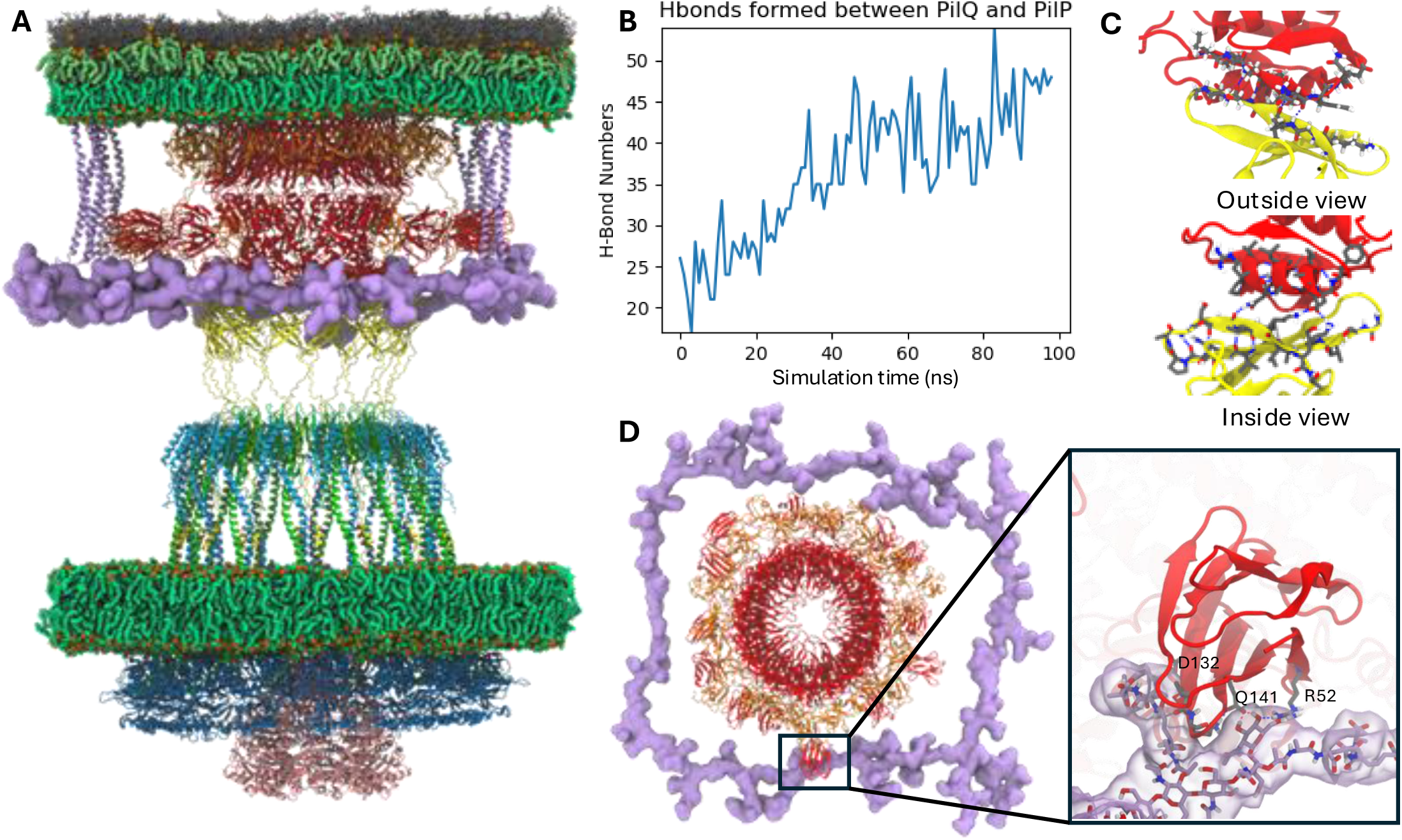
MD simulations of the CPM in its native environment. A) The all-atom system used for MD simulations. Proteins are shown in a cartoon representation with the following color scheme: PilQ in red, TsaP in orange, PilP in yellow, PilO in light blue, PilN in green, PilM in dark blue, PilT in pink, and Lpp in light purple. The glycopeptide is shown with a surface representation in light purple. Lipid atoms of the inner membrane and the inner leaflet of the outer membrane are represented as spheres, with carbon atoms in green and nitrogen in brown. Lipopolysaccharides (LPS) are illustrated in the outer leaflet of the outer membrane, where lipid atoms are shown as spheres with light green carbon and brown nitrogen atoms. The core and O-antigen regions of LPS are depicted in gray using a licorice representation. B) Number of hydrogen bond formed between PilQ and PilP during the simulation. C) Interactions between PilQ and PilP. The upper panel shows a view from the outside, while the lower panel shows a view from the inside. D) Top view of the system at 100 ns. Proteins and glycopeptide are represented in the same manner as in panel (A). An enlarged view highlights the interactions between PilQ AMIN domain and the glycopeptide. Important residues and the glycopeptide are shown in licorice representation, with glycopeptide carbons in purple, protein carbons in gray, oxygens in red, and nitrogen in blue. The glycopeptide surface is illustrated with transparent purple shading.

To assess the stability of interfaces between protein components, we analyzed hydrogen bond formation and contact areas between different CPM components (Fig. S20). A critical interface is between PilP and PilQ, connecting outer- and inner-membrane proteins, and we observed an increase in hydrogen bonds between these proteins (Fig. 5B). Detailed examination of a single PilQ-PilP interface showed the formation of hydrogen bonds on both the inner and outer sides (Fig. 5C). On the outer side, these interactions form a beta sheet, enhancing structural rigidity. On the inner side, we identified several salt bridges between an alpha helix and the beta sheet. Notably, these interactions were selective; not all protein pairs formed the same interactions (detailed hydrogen bonding changes between individual proteins are shown in Fig. S21).

Another interface of interest is found between PilQ and the PG (Fig. 5D), where an AMIN domain of PilQ forms stable hydrogen bonds with the PG. These interactions help to maintain the position of that AMIN domain during the simulation, suggesting that the PG contributes to stabilizing the PilQ-TsaP complex during operation.

### Dynamics of extension and retraction

We also conducted steered molecular dynamics (SMD) simulations to explore the pilus extension and retraction processes (Supplementary Movies 1 and 2). In these SMD simulations, the system focused on the upper half of the CPM, specifically the OM portion including PilQ and TsaP, coupled with the PG and a short pilus comprising five major pilins (PilA). By applying SMD, we pulled this pilus fragment through the secretin (PilQ) from the periplasmic side to the extracellular side to mimic the extension process. Based on the structure obtained from the extension SMD simulations, we applied force to draw the pilus fragment from the extracellular side back into the periplasm to mimic the retraction process.

We performed three replicates for each process at a pulling speed of 5 Å/ns. In extension simulations, peak forces around 5,000 pN were required when the largest cross-sectional area of the pilus passed through the channel (Fig. 6A). Retraction simulations showed peak forces near 4,000 pN, fluctuating during major pilin passage (Fig. 6B). Although the peak forces are much higher than those used *in vivo* due to the pulling speed, qualitative observations are still relevant^49^. The PilQ channel features a gate formed by two layers of beta strand loops. In the extension SMD simulations, this gate is opened as the major pilin subunits push against these loops (Fig. 6D). The tips of the beta strand loops form and maintain interactions with the pilin subunits, while other parts of the loops partially maintain the hydrogen bonds that stabilize their beta strand structure. During retraction, the gate is initially open toward the extracellular side. The tips of the loops are pushed back by the retracting pilus, but most of the loops remain oriented outward (Fig. 6E). Thus, these dynamic loops seal the channel around the pilus, allowing only the pilus to pass through during both the extension and retraction processes.

**Figure 6.**
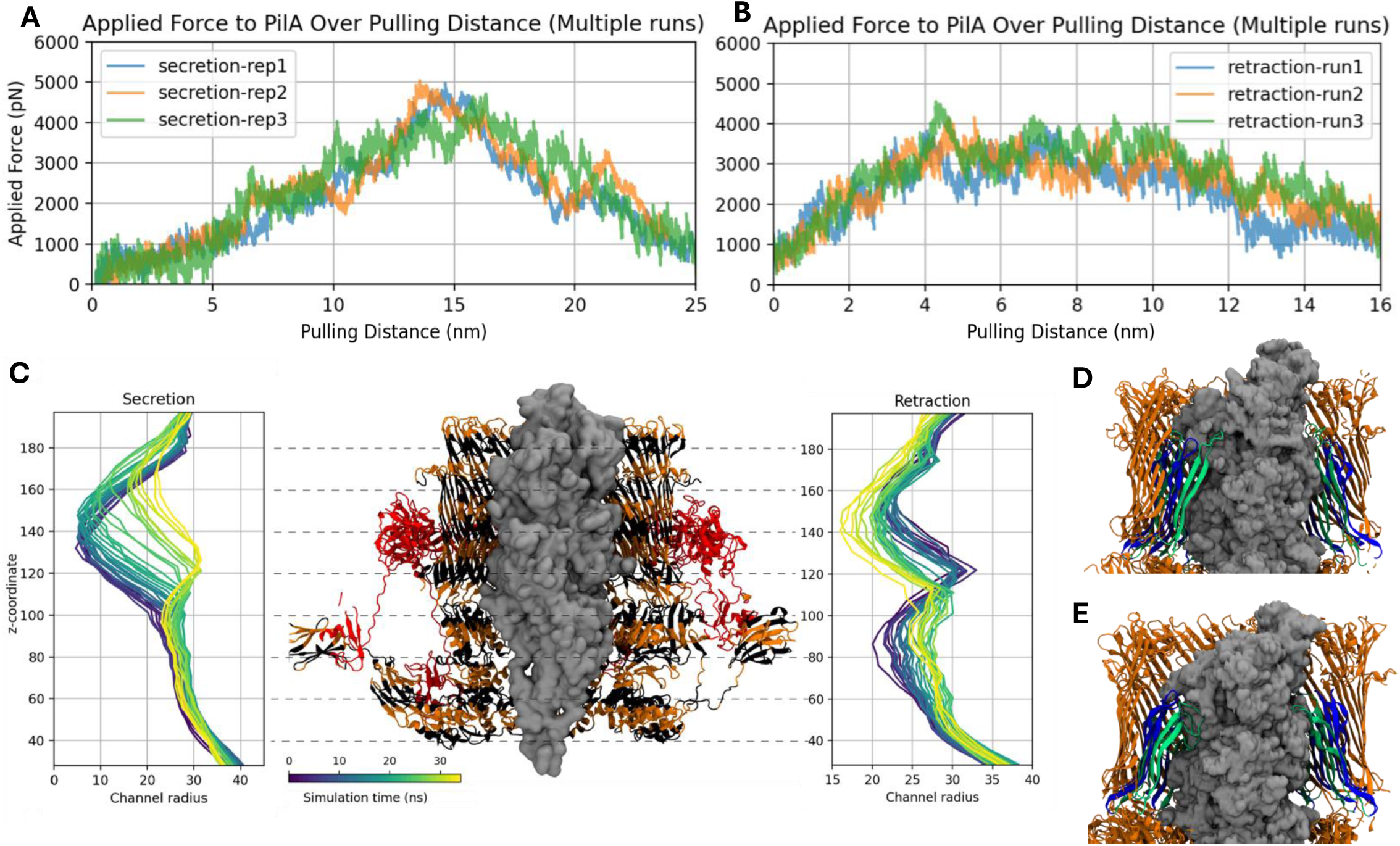
SMD simulations of PilQ, TsaP, and PilA in their native environment. A) Force-distance profiles for all three replications of the secretion simulation. B) Force-distance profiles for all three replications of the retraction simulation. C) Changes in the channel radius of PilQ during the simulations: the left panel illustrates the secretion simulation, the right panel shows the retraction simulation, and the middle panel indicates the corresponding positions along the z-axis. D) Snapshot from the secretion simulation showing the open conformation of the PilQ gate, with the two layers of beta strand loops highlighted in green and blue. E) Snapshot from the retraction simulation displaying the open conformation of the PilQ gate, with two layers of beta strand loops highlighted in green and blue.

During extension, PilQ displayed remarkable flexibility, with the channel radius growing from 5 Å to 32 Å at the gate region. For retraction, the radius varied from 33 Å to ∼20 Å (Fig. 6C). For the extension and retraction processes, the maximum channel radius did not differ significantly between the two, indicating that the PilQ channel maintained its integrity during pilus translocation. This balance between flexibility and stability likely plays an important role in accommodating the passage of the pilus while maintaining the structural framework of the CPM.

## Discussion

In this work we investigated the CPM from multiple perspectives, ranging from the quasi-atomic resolution of the fiber to the nanoscale *in-situ* architecture of the machine, which allowed us to build a complete pseudoatomic integrative model. We then used the model to simulate the retraction-extension cycle, which provided insights into the dynamics and mechanical properties of the pilus.

The cryo-EM structure of the filament and functional experiments allowed us to identify residues involved in filament auto-aggregation (PilA^K40^, PilA^K108^). These residues are surface-localized and contribute to the network of positively and negatively charged patches along the helical symmetry of the pilus. Auto-aggregation is observed in several T4a filaments and is involved in adhesion and micro-colony formation, and it also enhances plasmid exchange in liquid^50^. Inter-filament interactions are usually modulated by post-transcriptional modification^51,52^, indicating the charge-mediated aggregation of the CP might be unique in the type IVa family.

Cryo-ET investigation of the *in-situ* CPM architecture in the Δ*pilT* strain revealed new features of its non-piliated and piliated states. Both piliated and non-piliated STAs of the Δ*pilT* strain show no cytoplasmic density, likely due to low PilB occupancy, which is in line with real-time fluorescence microscopy showing brief bursts of extension^7^. We identified two classes of the non-piliated state showing the absence or presence of a lower periplasmic ring, but, given the static nature of cryo-ET, it is not clear whether the absence of this ring is due to assembly or disassembly. Similarly, the piliated state also showed two classes with a difference in the distance between the mid-and lower periplasmic rings. We could not detect machines with an intermediate distance between these rings, leading us to believe these classes are two discrete states. The compacted state could be the result of the slow PilT-independent pilus retraction. PilA disassembly from the pilus fiber occurs in the outer leaflet of the IM, where the PilQ gate might act as a clamp, resisting fiber sliding and causing the periplasm to shift locally. This would slowly draw the mid-periplasmic ring closer to the OM pore density until steric clashes occur^19^. Further investigation will be needed to completely understand these two states.

A second new feature of the CPM is the presence of a ring flanking the secretin density. By imaging a PilQ-GFP strain, we identified this region as the PilQ AMIN domain. In the single particle reconstruction map of PilQ, this domain was absent, probably due to high flexibility of the structure^21^. *In situ*, this domain anchors PilQ to the PG. Because *Vc*PilQ has only one AMIN copy per protein, the position of this domain would consistently be around the ring, producing the observed flanking density.

The CP utilizes two retraction ATPases, PilT and PilU. Our cryo-ET data clearly shows that PilU forms an additional ring bound to PilT. Because both PilT and PilU form independent homohexameric rings^17^, the PilT-PilU assembly is formed by a dimer of hexamers. PilU can partially restore the retraction abilities of nonfunctional PilT^17^, indicating that direct PilU binding might boost PilT processivity or allosterically induce PilT rearrangements that lead to forceful retraction.

We took advantage of the extended literature on type IVa pili and the advent of AI-driven structural protein predictors to build a complete pseudoatomic model of the CPM. According to our model, the space between the mid- and lower periplasmic rings that undergoes compaction is occupied by an unstructured region in PilP (residues 66–92) that could allow this rearrangement. Our model also suggests that PilB and PilT might contact PilC in opposite orientations. PilB and PilT are homologous proteins that each contain Walker A and Walker B motifs. In the current model, these proteins hydrolyze ATP, inducing conformational changes in the ATPase ring that force PilC to rotate either clockwise (leading to pilus extension) or counterclockwise (leading to pilus retraction)^42,53,54^, but the exact mechanism of this process is still unknown. Our model of the IM platform suggests that the opposite PilC rotations induced by PilB and PilT occur because each ATPase binds to PilC upside down in relation to its homolog. The N-terminal domain of PilB binds PilC and hydrolyzes ATP normally, rotating PilC clockwise, while PilT is flipped upside down, using its C-terminal domain to bind PilC. It hydrolyzes ATP in the same direction as PilB, but because it is upside down, it rotates PilC in a counterclockwise direction. This differential binding might explain how a single ATPase is able to catalyze both extension and retraction in Archaeal and tight-adherence pilus systems: If a single ATPase is able to mediate pilus extension, then flip upside down and mediate retraction, it would have no need for a second retraction-focused ATPase.

Our MD simulations demonstrated the structural stability and dynamics of the CPM model in a near-native environment, highlighting its robustness under physiological conditions. The 120-ns MD simulations revealed stable interactions between key components, including PilP-PilQ and PilQ-PG, which were reinforced by hydrogen bonding and salt bridge formation. Steered MD simulations of pilus extension and retraction further revealed how the structure of the secretin supports an impressive interplay between flexibility in the gate, which seals the channel until the pilus passes through, and stability in the beta barrel, which remains intact throughout. The secretin’s domains appear efficient and minimized for their purposes—the gate is just a ring of thin, flexible, interacting loops, and the beta barrel is just one layer thick. We wondered whether the pitch of the beta strands might change in response to the pilus protruding (like a Chinese finger trap), but they do not—the barrel is just the right size to accommodate the open gate and pilus. The tight fit of the pilus within the secretin indicates that DNA is taken up at the pilus tip rather than entering orthogonally, consistent with fluorescence data showing DNA binding at the tip^7^. These findings suggest that CPM assembly is structurally stable while staying functionally malleable to ensure efficient pilus translocation, and together they exemplify the idea of structure determining function.

## Materials and Methods

### Cell growing condition and cryo-ET sample preparation

*Vibrio cholerae* strains were grown in LB, 100 μg/mL carbenicillin, 50 μg/mL kanamycin, 25 μg/mL zeocin in a shake incubator at 30°C. Following overnight growth, 50 μL of the culture was spread on LB agar plates containing 10 mM CaCl2, 20 mM MgCl2, 0.1 mM IPTG and incubated for 48 h at 30°C. Cells were collected from plates using cotton swabs, which were then dipped into Eppendorf tubes containing 7 g/L aquarium salt (Instant Ocean Sea Salt) and gently rotated to resuspend the bacteria in solution. The suspension was centrifuged at 10,000 x g (3 min) and the pellet was washed once with aquarium salt solution, followed by centrifugation.

### Pilus purification

Pili were shed from bacteria by resuspension of the washed pellet in 50 mM Tris-HCl, pH 9.5, followed by vortexing (1 min). Cells were removed by four spins at 6,000 (3 min), after which the supernatant was transferred to new tubes and spun again.

The subsequent procedure takes advantage of the tendency of competence pili to form dense bundles at pH values <10 and dissociate into single filaments at pH >10. The sample was first dialyzed for 4 hours at 4°C against 50 mM Tris-HCl, pH 7.4, using a 300 MWCO dialysis bag (Spectra/Por, Biotech). It was then centrifuged at 15,000 x g (1 h) to remove DNA and small contaminants. The pellets (bundled pili) were resuspended in 400 μL 50 mM Tris-HCl, pH 7.4, and dialyzed against 150 mM Tris-HCl, 100 mg Lysine, pH 8.5 for 6–8 hours. The high MWCO removes mid-sized contaminants, and the lysine destabilizes pili bundles so that they can fully dissociate in the subsequent step. The bag was transferred to 150 mM ethanolamine, pH 10.6, and dialyzed overnight. The dialyzed sample was centrifuged at 10,000 (1 h) and the supernatant (dissociated pili) were concentrated using a Vivaspin® 30 MWCO to a concentration of 1 mg/mL. The presence of intact pili was verified by negative staining with 3% uranyl acetate on 300 mesh formvar/carbon grids (Electron Microscopy Sciences) using a 120 kV FEI Ti12 with LAB6 filament and Gatan Ultrascane 2k x 2k CCD camera.

### Cryo-EM and cryo-ET sample preparation and data collection

The pili were cryogenically frozen using a FEI Vitrobot MK4 on glow-discharged R1.2/1.3, 300 mesh UltrAufoils EM-grids (Electron Microscopy Sciences) at 22°C and 100% humidity (blot time of 6 s; blot force of 5 s; with 3 s wait time). For cryo-ET the cell suspension was concentrated by centrifugation to 10 mg/mL (bacteria dry weight/aquarium salt solution) and 100 μL of this solution was pelleted by centrifugation. The cell pellet was mixed with 20 μL of 10-nm gold colloidal beads (Sigma-Aldrich) pretreated with BSA (5% in PBS). R2/2 carbon-coated 200 mesh copper Quantifoil grids (Quantifoil Micro Tools, GmbH, Jena, Germany) were glow discharged for 60 seconds to prepare hydrophilic surfaces before sample application. A 3 μL volume of the cell and gold beads solution was deposited on to the EM grids and plunge-frozen using a FEI Vitrobot MK4 at 22°C with 100% humidity (blot time of 8 s; blot force of 3 s; with 10 s wait time).

Data collection was done at the Beckman Institute Resource Center for Transmission Electron Microscopy at Caltech using a 300 kEV FEI Titan Krios equipped with a Gatan K3 director and Gatan energy filter. The Serial-EM software^55^ was used for data acquisition. Pili micrographs were acquired at a defocus range of -0.6 to -2 and calibrated pixel size of 0.416 Å, binned to 0.832 Å during collection. Movies of 40 frames were collected with a total dose of 60 e^-^/Å^2^.

Tilt series were acquired using FISE pipeline, tilting the sample from -60 to 60 degrees with 3-degree increments. A total dose of 160 e^-^/A^2^ was applied with a defocus of -8 um, pixel size of 3.3 A, and magnification of 26,000X.

### Helical Reconstruction

The data was processed within the RELION framework by helical reconstruction^27,56^. Motioncorr2^57^ and CTFFIND-4.1^58^ were used for motion correction and CTF-estimation, respectively. Following manual particle picking in which the start and end coordinates of pilus filaments were defined, 300-px boxes were extracted with a 10 Å overlap (estimated to correspond to a helical rise). 77,000 overlapping boxes were selected following 2D classification. A cylinder with a diameter of 80 Å was used as a starting reference for the initial 3D classification, and it was processed without applying helical symmetry to avoid bias from incorrect helical parameters. The highest resolving 3D class average was selected, and the helical parameters of this reconstruction were extracted using the RELION plugin Relion_Helix_Toolbox. The original 77,000 boxes were then subjected to two rounds of 3D classification with the estimated starting helical parameters and C1 symmetry. The helical parameters were refined within a specific range (for the final dataset: rise: 10.6 -11.2, twist: 92.2 -94.8. Larger ranges were allowed in preliminary processing to optimize refinement settings). We selected 22,000 particles following 3D classification and refined them using the central 60% of the filament to avoid Fourier artifacts at the edges using a mask extending 6 px that was generated in RELION. The resolution following CTF refinement, Bayesian polishing, and a last round of 3D refinement resulted in a map with an estimated resolution of 3.3 Å as defined by FSC.

### Subtomogram averaging and difference analysis

Competence pilus machines were identified by visual inspection. The IMOD package was used to create three-dimensional reconstructions of tilt series^59–61^ and subtomogram averaging. PCA and K-means clustering were performed using the PEET sub-volume averaging program with two-fold symmetrization along the y-axis^30,31^. For the difference analysis all the particles were re-aligned to the ΔPilT STA and the central slice image was imported in GIMP for subtraction.

### Model Building

The PilA starting model was generated by homology modeling using SWISS-MODEL^62,63^ and the *P. aeruginosa* PAL pilin (1OQW) as an input model. The density map was sharpened in Phenix^64^ using the two unmasked half-maps and segmented into densities corresponding to pilin subunits using the Chimera plugin Segger v1.9.5^65,66^. A single pilin subunit was first refined into the cropped map density corresponding to a single subunit using Phenix Refine and Coot^67^. When a good fit was achieved, 14 total refined copies were docked into the full density map and further refined in Phenix with NCS constraints enabled. This model was then docked into the single segment map and refined in Phenix and Coot. This process was repeated until refinement did not improve the qualitative model parameters in the full, multi-subunit model.

For a complete description of how we built the CP machine model, see the Supplementary Information.

Briefly, we used solved structures as a starting point to dock AlphaFold2^36^ and ColabFold^37^ models. We started from the *Vc*PilQ solved structure and used the solved structures of PilQ-PilP (PDB:4AV2)^68^ and PilQ-TsaP (6VE2)^22^ to position the *Vc*PilP and *Vc*TsaP models (Fig. S7–8). In the PilQ-PilP case we further refined the docking using HADDOCK and conserved residues in the interacting interface as anchor points (Fig. S9).

We modeled PilN_1_PilO_1_PilP_1_^69^, PilN_1_PilM_1_PilB_1_^45^, PilC_2_PilB_6_^19,28^, PilC_2_PilT_6_, and PilU_6_^17^ according to literature using ColabFold multimer. We checked the quality of the models by using information from experimentally characterized homologous systems. All the models were fitted in their respective densities and symmetrized to match the PilQ solved structure.

### System construction for MD simulations

The atomistic model described above was placed into a dual-membrane environment for MD simulation. We constructed the inner membrane (IM) and outer membrane (OM) using CHARMM-GUI^70^ and equilibrated them separately. The IM was built around PilC, PilO, and PilN atop PilM, while the OM was constructed around PilQ and TsaP. Specifically, lipopolysaccharide (LPS) from *Vibrio cholerae* serotype O1 was used for the outer leaflet of the OM^71^. The inner leaflet of the OM, as well as both leaflets of the IM, consisted of 4% POCL2, 10% POPG, and 86% POPE^72^. Both membranes were built to approximately 295 Å × 295 Å within a water box, and 0.15 M KCl was added to the solution. The OM-protein and IM-protein systems were released and equilibrium step by step separately^73^, followed by simulations with the x-y dimensions constrained to maintain a constant ratio until the membrane dimensions stabilized.

The PG cell wall (CW) model was constructed by reverse-coarse-graining a PG patch under tension from previously published simulations of a coarse-grained sacculus^74^. The patch was selected to have a sufficiently large opening to accommodate the CPM. The CW was connected to the OM through four lipoprotein (Lpp) trimers, and AlphaFold2^36^ was used to predict the Lpp structure based on the sequence (Uniprot: A0A085PVQ9).

After building and equilibrating each part separately, we combined them to form a comprehensive complex system that includes the IM, OM, CW, CPM, water, and ions. Trial simulations were conducted to determine the amount of water in the periplasm necessary to achieve a density of 1 g/cm^3^ and to avoid compression or expansion. The final system comprised around 4.9 million atoms, with the size of 300 Å × 300 Å × 533 Å. Additionally, we built a reduced system for steered MD simulations, which contained the OM, TsaP, PilQ, Lpp, CW, and a fragment of the major pilin, along with water and ions. The major pilin fragment was constructed using 10 PilA monomers, equilibrated separately, while all other components were derived from the OM protein systems described earlier. This reduced system consisted of approximately 4.5 million atoms, with dimensions of 300 Å × 300 Å × 490 Å.

### Simulation methods for MD simulation

All-atom MD simulations were performed using NAMD2.13^75^ and NAMD3^76^ with the CHARMM36m force field parameters for proteins^77,78^, CHARMM36 parameters for lipids^79^, and the TIP3P-CHARMM water model^80^. A previously developed PG force field was also used^81^. All simulations were carried out under periodic boundary conditions with a cutoff at 12 Å for short-range electrostatic and Lennard-Jones interactions and a force-based switching function starting at 10 Å. Long-range electrostatic interactions were calculated using the particle-mesh Ewald method^82^ with a grid spacing of 1 Å. Each system was equilibrated under an isothermal-isobaric ensemble (NPT) at 310 K and 1 atm with a timestep of 4 fs utilizing hydrogen mass repartitioning^83,84^. Temperature control was maintained using a Langevin thermostat with a damping coefficient of 1 ps⁻¹, and pressure control was achieved using a Langevin piston^85^. VMD was used for all visualization^86^.

### Simulation methods for Steered MD

SMD simulations were conducted using the collective variables module (Colvars)^87^, focusing on the OM-protein part of the system. Force was applied to every 10th C*_α_* atom of the major pilin fragment, in the +z direction to mimic extension and in the −z direction to mimic retraction along the membrane normal. The pulling speed was set to 5 Å/ns, and the force constant was 10 kcal/mol/Å^2^. To maintain the shape and orientation of the major pilin fragment, orientation and root-mean-square deviation (RMSD) restraints were applied during the SMD simulations. To stabilize the position of PilQ during the simulations, restraints were applied to the following two distances: 1) that between the top residues of PilQ and the lipid A headgroups of the LPS in the outer leaflet of the OM and 2) that between the residues at the interface of PilQ and the inner leaflet lipid headgroups with the inner leaflet lipid headgroups themselves.

These restraints ensured that PilQ maintained its position relative to the OM throughout the simulations. The extension simulations required ∼55 ns, and the retraction simulations required ∼35 ns. Each system’s SMD production run was performed three times.

### Chitin-independent natural transformation assay

For induction of competence in *V. cholerae*, the master competence regulator, Tfox, was placed under an IPTG-inducible promoter. Cells were also genetically locked in a high cell density state by deletion of *luxO*. To induce cells, strains were grown to late-log in LB supplemented with 20 mM MgCl_2_, 10 mM CaCl_2_, and 100 µM IPTG by rolling at 30 °C. Then, 7 µL of this culture was diluted into 350 µL of instant ocean medium (IO; 7g/L Aquarium Systems). Two transformation reactions were setup for each strain: one where 100 ng of transforming DNA (ΔVC1807::Kan^R^ tDNA) was added and a negative control where no tDNA was added. Reactions were then incubated statically at 30 °C overnight. The next day, 0.5 mL LB was added to each reaction and transformations were outgrown at 37 °C shaking for 3 hrs. Reactions were then serially diluted and dribble plated onto LB + Kan50 (to quantify transformant CFUs) and plain LB agar plates (to quantify total CFUs). The transformation frequency was determined by dividing the transformant CFUs by the total CFUs. For samples with no transformants, the limit of detection was calculated and plotted.

### Pilus labeling and imaging

Strains were grown exactly as described above for chitin-independent natural transformation assays. Then, 100 µL of late-log cells were pelleted at 8000 x g for 1 min and washed in IO. Cells were then labeled with 25 µg/mL AF488-mal for 15 mins static at room temperature. Cells were then washed three times in IO, ensuring to spin at 8000 x g for 1 min to prevent shearing of surface pili. Next, 2 µL labeled cells were placed under an 0.4% IO gelzan pad and imaged with the phase and FITC channels on an inverted Nikon Ti-2 microscope with a Plan Apo ×60 objective fitted with a Hamamatsu ORCAFlash 4.0 camera using Nikon NIS Elements imaging software. Representative images of the piliation phenotype for each strain were selected from the raw microscopy images.

### Cellular aggregation assay

Strains were grown exactly as described above for chitin-independent natural transformation assays. Cultures were allowed to sit static at room temperature on the benchtop for 15 mins to allow aggregated cells to settle out of solution. Representative images were taken of each strain without disturbing settled cells.

## Supporting information

Supplemental Modeling Information

Supplementary Movie 1: Extension

Supplementary Movie 2: Retraction

## Acknowledgements

This work was supported in part by the National Institutes of Health (R01 GM148586 to JCG, R35 GM128674 to ABD, and RO1 AI127401 to GJJ). SK was supported by the Swedish Research Council (2019-06293). Cryo-electron microscopy was performed in the Beckman Institute Resource Center for Transmission Electron Microscopy at Caltech, and we thank Songye Chen for her help coordinating use of the microscope. MD simulations were run using Summit, a resource of the Oak Ridge Leadership Computing Facility at the Oak Ridge National Laboratory, which is supported by the Office of Science of the U.S. Department of Energy under Contract No. DE-AC05-00OR22725.

## Author contributions

SM, SK, ABD, JCG, and GJJ conceived the project; SK purified the competence pilus, performed cryo-EM, and built the competence pilus model; SM performed bioinformatics, cryo-ET, sub-tomogram averaging, and modeling of the competence pilus machine; LY and DLL ran MD simulations; AET performed transformation and aggregation assays. SM wrote the manuscript with support from all authors. ABD, JCG, and GJJ provided training, supervision, and resources throughout the project.

---

**Figure S1.**
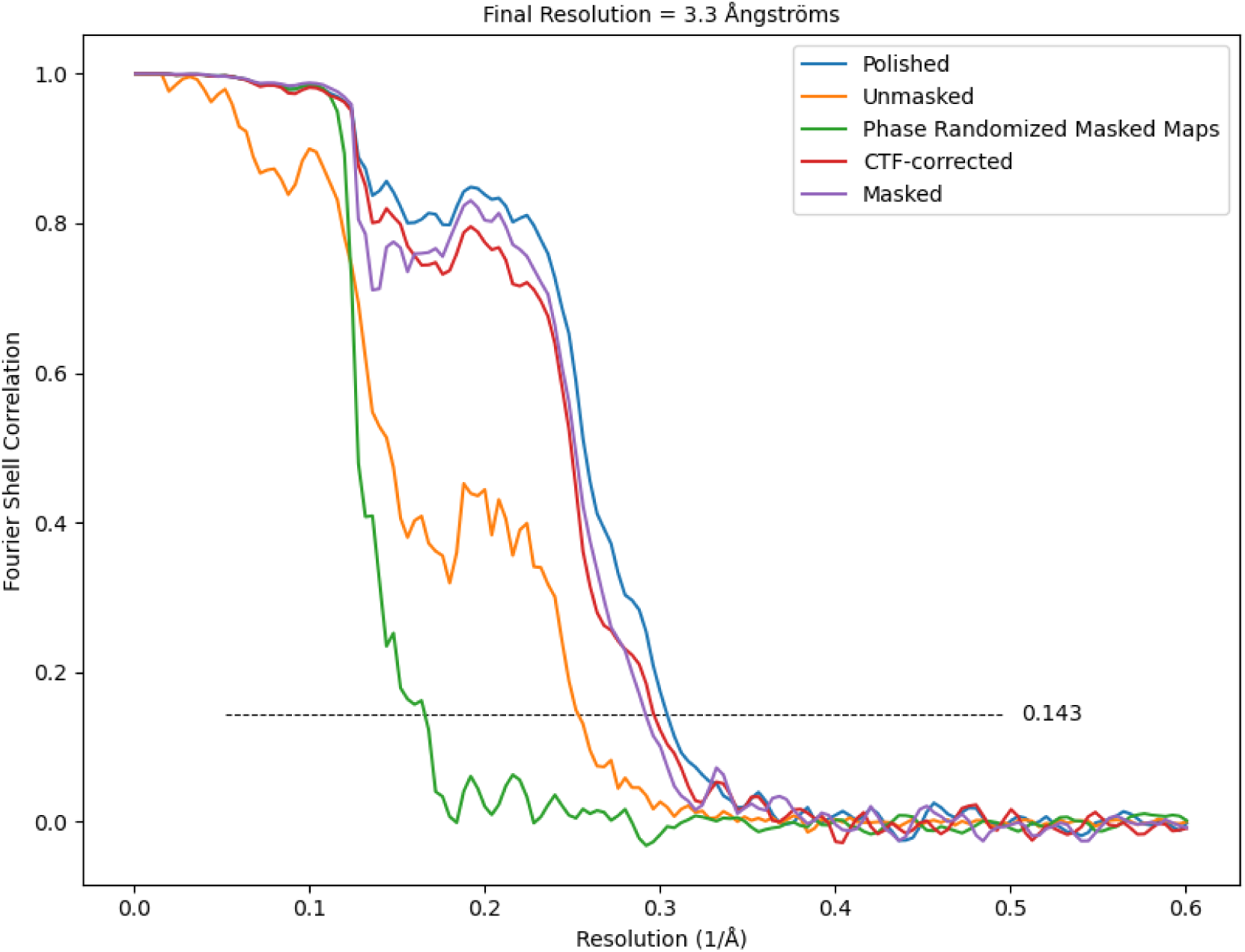
CP Resolution. Relion’s FSC curves of the competence pilus helical reconstruction showing a final resolution of 3.3 Å.

**Figure S2.**
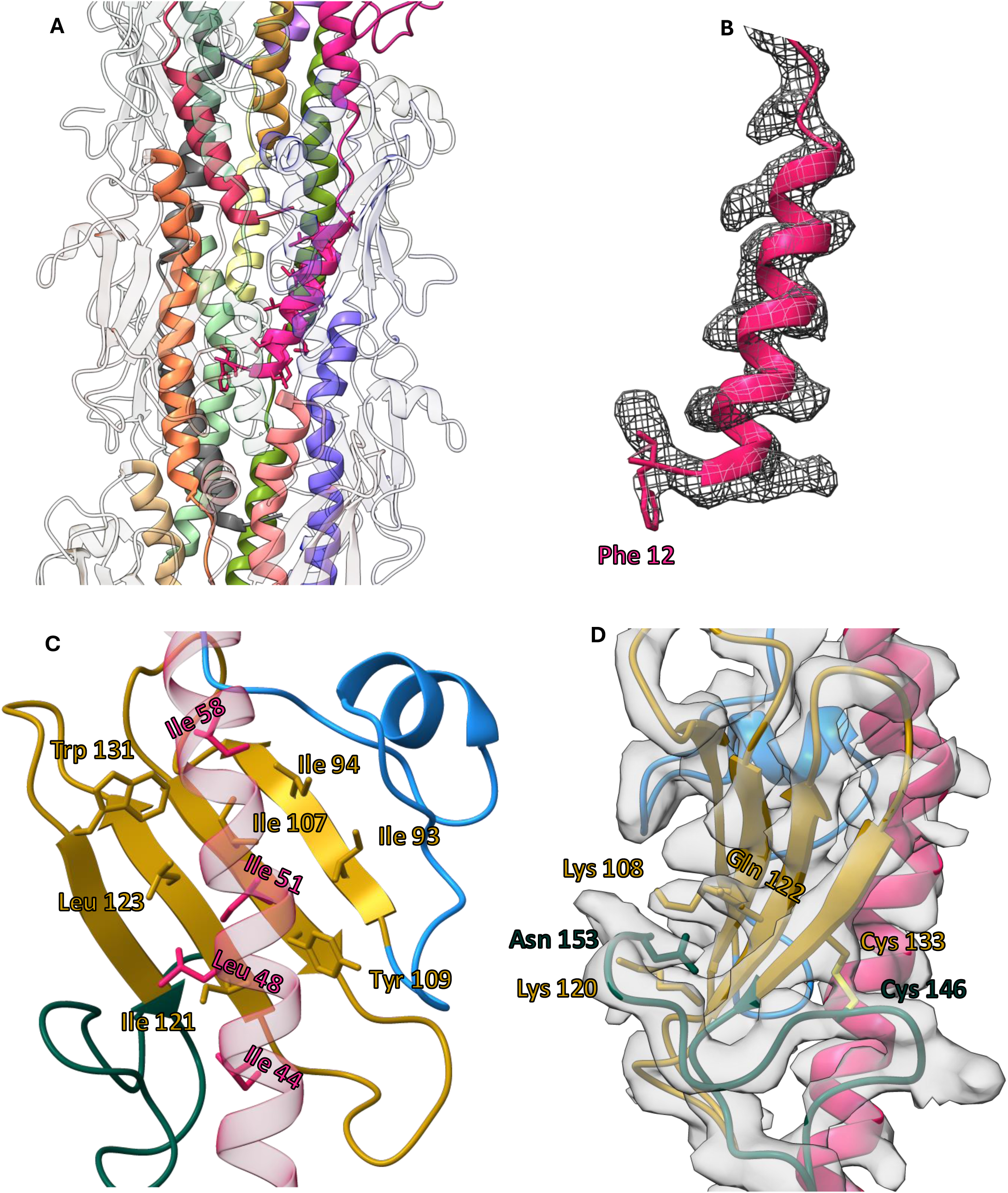
PilA structure. A) The N-terminal domain of PilA is buried inside the CP filament and is stabilized by hydrophobic residues (shown in stick from position 12: FTLIELMIVVAVIGVLAA) B) Structure of the methylated phenylalanine (methyl group up, phenyl-group down) and the respective cryo-EM map. C) Highlight of the beta sheet domain with hydrophobic residues shown in stick. D) Stabilization of the C-terminal domain by the C133-C146 disulfide bond and by possible electrostatic interaction between Asn 154 and nearby charged residues (Lys 108,120, and Gln 122 )

**Figure S3:**
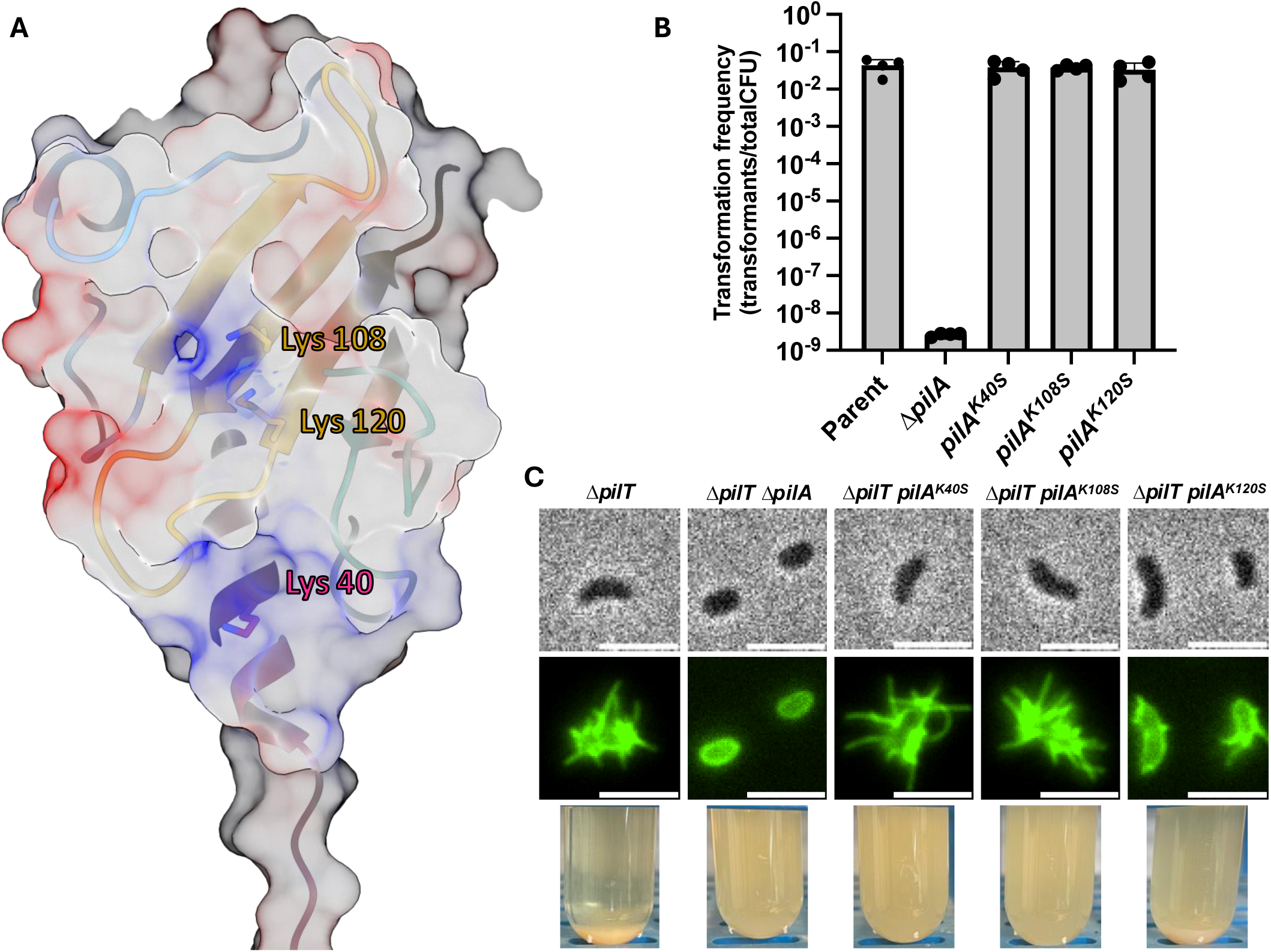
PilA lysines 40 and 108 are involved in inter-pili interaction. A) Localization of PilA^K40^, PilA^K108^, and PilA^K120^, which form the positively charged patches on the pilus surface. B) Transformation frequency of the parent, *ΔpilA*, and PilA surface-exposed lysine to serine substitutions (*pilA*_K40S_*, pilA*_K108S_, and *pilA*_K120S_). C) Phase contrast (top), fluorescence images (middle) of *Vibrio cholerae* with PilA stained with 488-malemide (scale bars 3 µm) and flocking assay (bottom) of *ΔpilT*, *ΔpilT ΔpilA*, and PilA surface-exposed lysine to serine substitutions in a *ΔpilT* background (*pilA*_K40S_*, pilA*_K108S_, and *pilA*_K120S_). Only the *pilA*_K40S_ and *pilA*_K108S_ substitutions prevent cell flocking without affecting transformation efficiency and pilus assembly.

**Figure S4.**
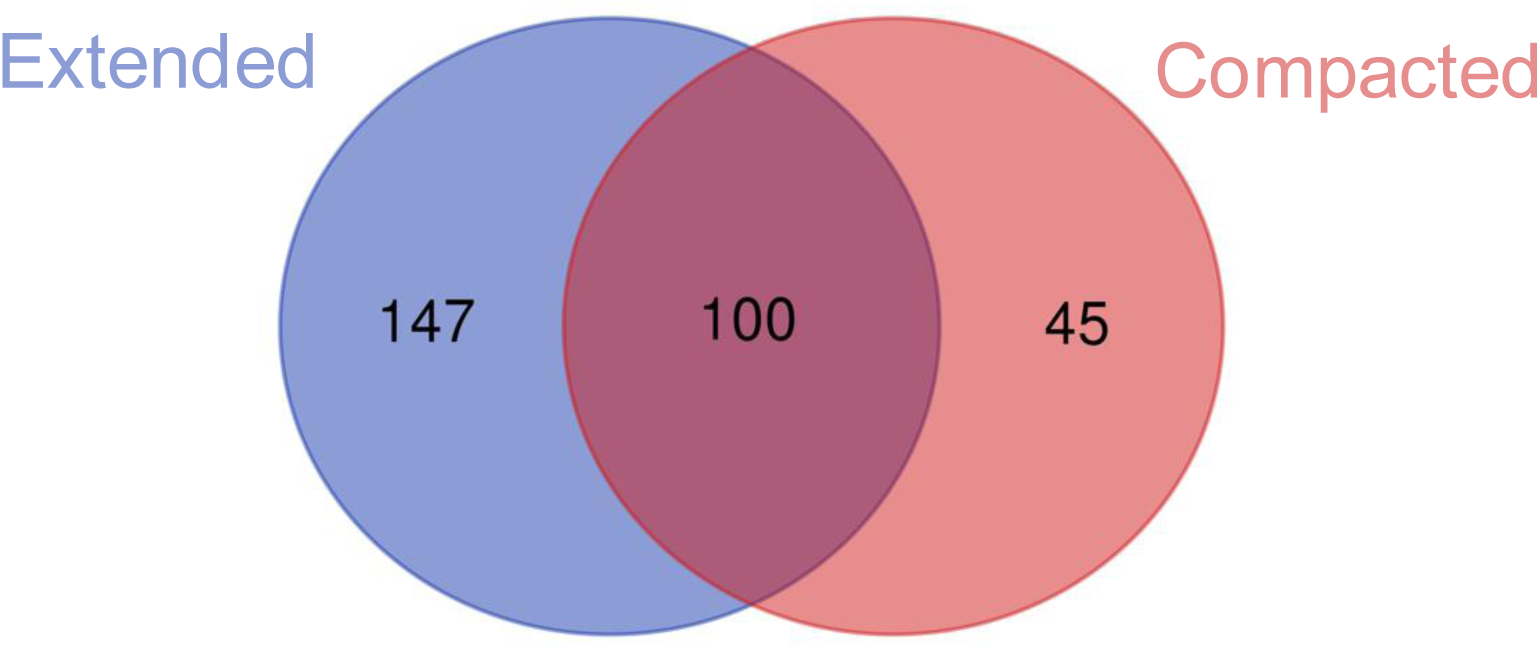
The compacted and the extended states coexist in *Vibrio cholerae* cells. After classification of the compacted and extended states of the CPM, we looked for tomograms that contain both states. Of 292 tomograms, 147 contain only the extended state, 45 have only the compacted state, and 100 contain a coexistence of the two states.

**Figure S5.**
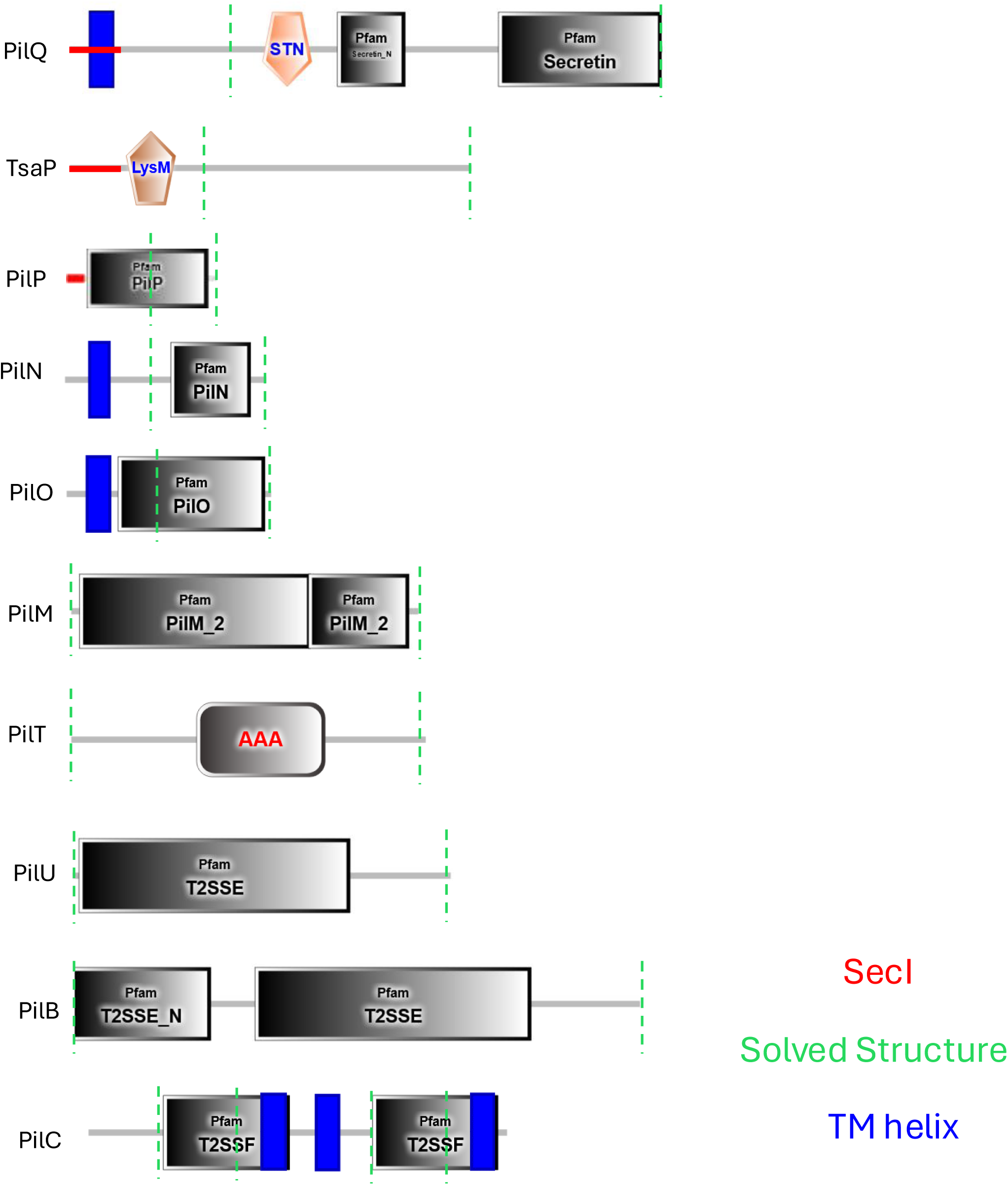
Domain architecture of the proteins involved in the assembly of the CPM. Red shows the presence of a SecI signal peptide, blue shows the presence of a transmembrane helix, and green highlights the portion of proteins whose structure has been solved (homologs included).

**Figure S6.**
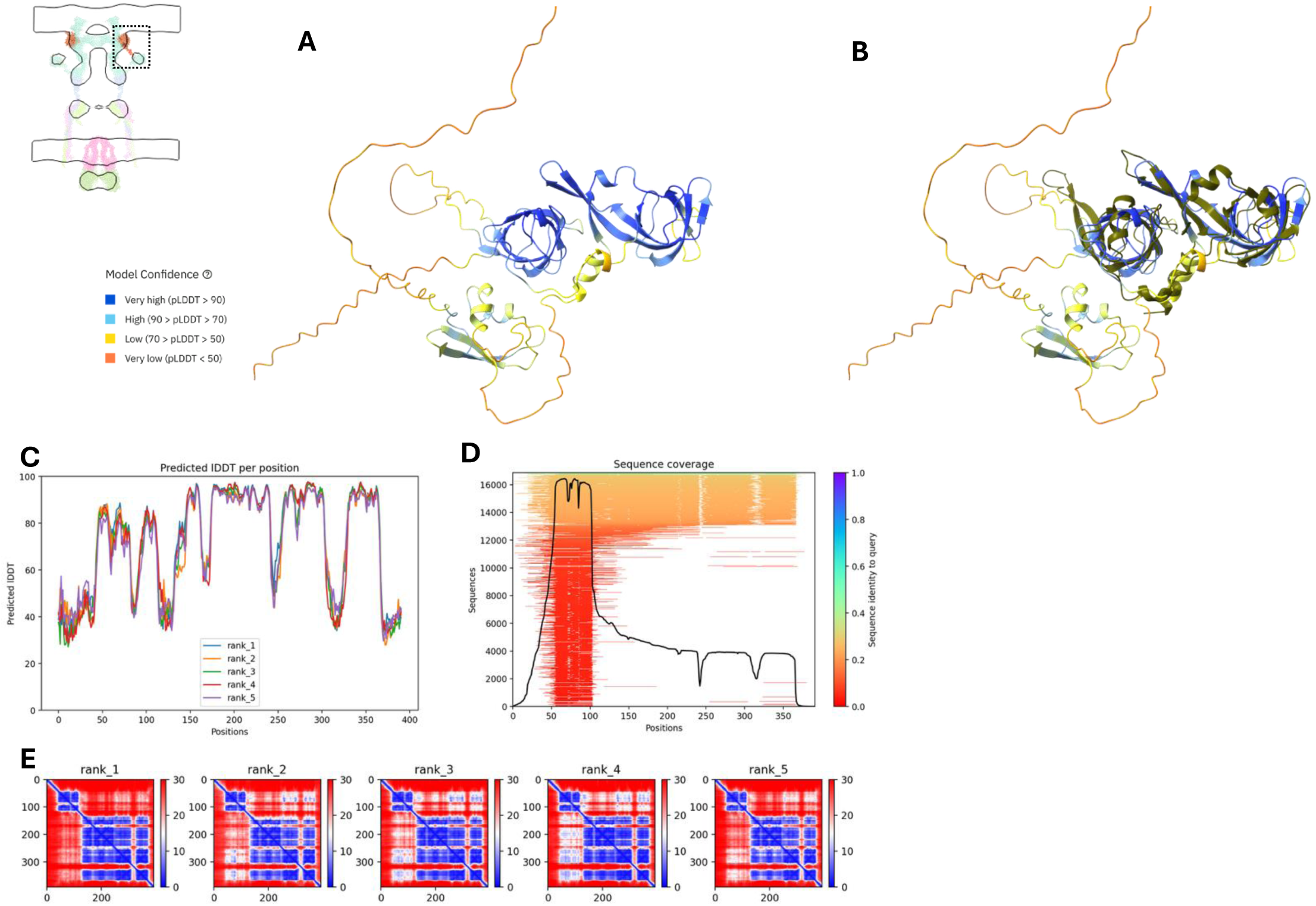
TsaP AlphaFold (AF) model. A) AF model of TsaP shows a bimodular organization. The N-terminus contains the LysM domain, which is responsible for PG binging. The C-terminus has a higher model confidence and B) superimposes well with the solved structure from *Pseudomonas aeruginosa* (6VE2) (RMSD between 71 pruned atom pairs is 1.3 Å; across all 206 pairs is 5.6 Å). C) AF predicted local Distance Difference Test, D) Sequence coverage and E) Predicted Aligned Error matrix (colormap in Ångströms).

**Figure S7.**
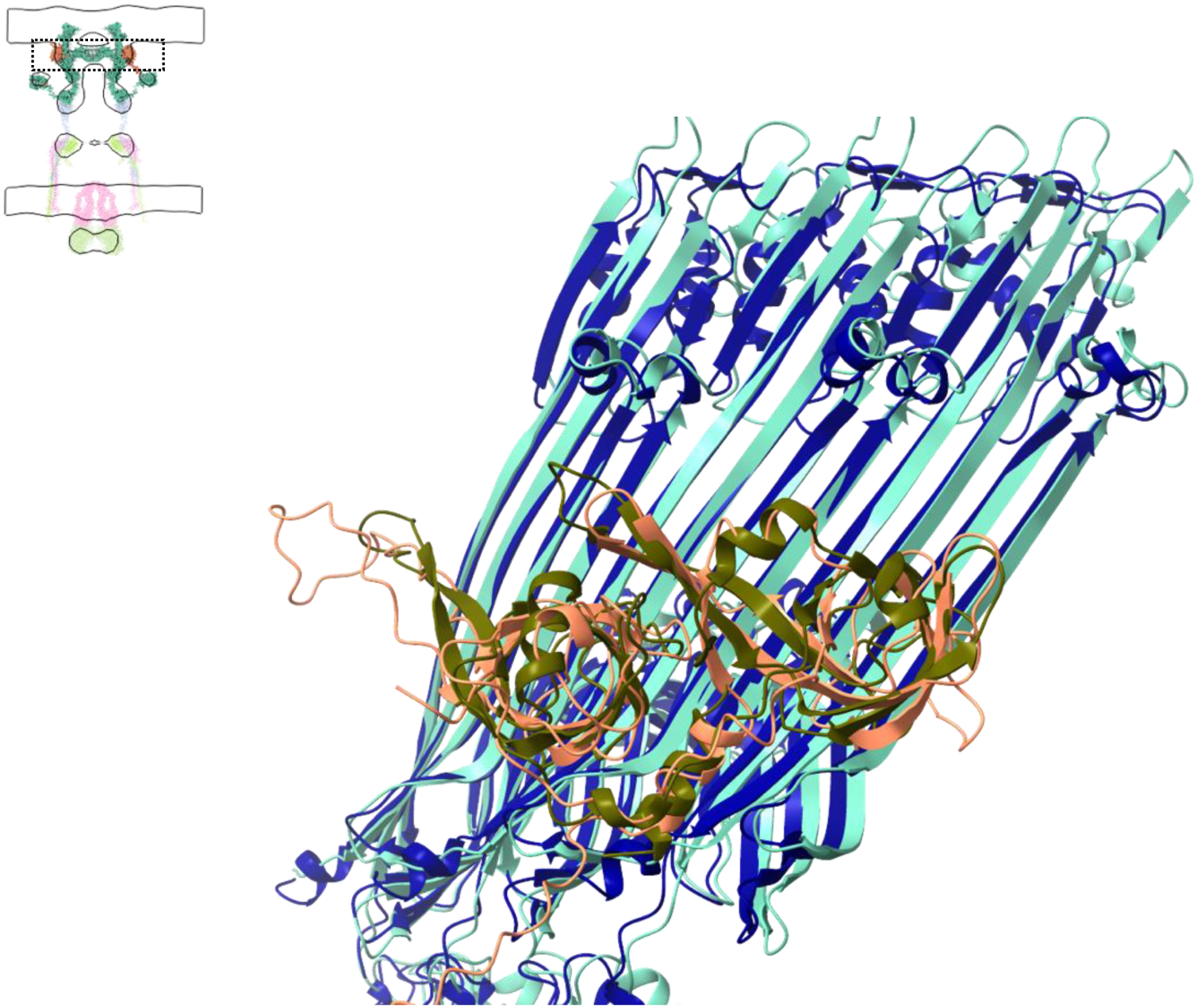
TsaP AF docking on VcPilQ using the *Pseudomonas aeruginosa* structure as a scaffold. Given the good alignment of *Vc*TsaP and *Pa*TsaP, we used the *P. aeruginosa* solved structure as a scaffold to dock *Vc*TsaP to *Vc*PilQ . The *Vc*PilQ-*Pa*PilQ alignment score is 994.1 with a RMSD between 142 pruned atoms of 1.1 Å; across all 294 pairs is 7.2 Å.

**Figure S8.**
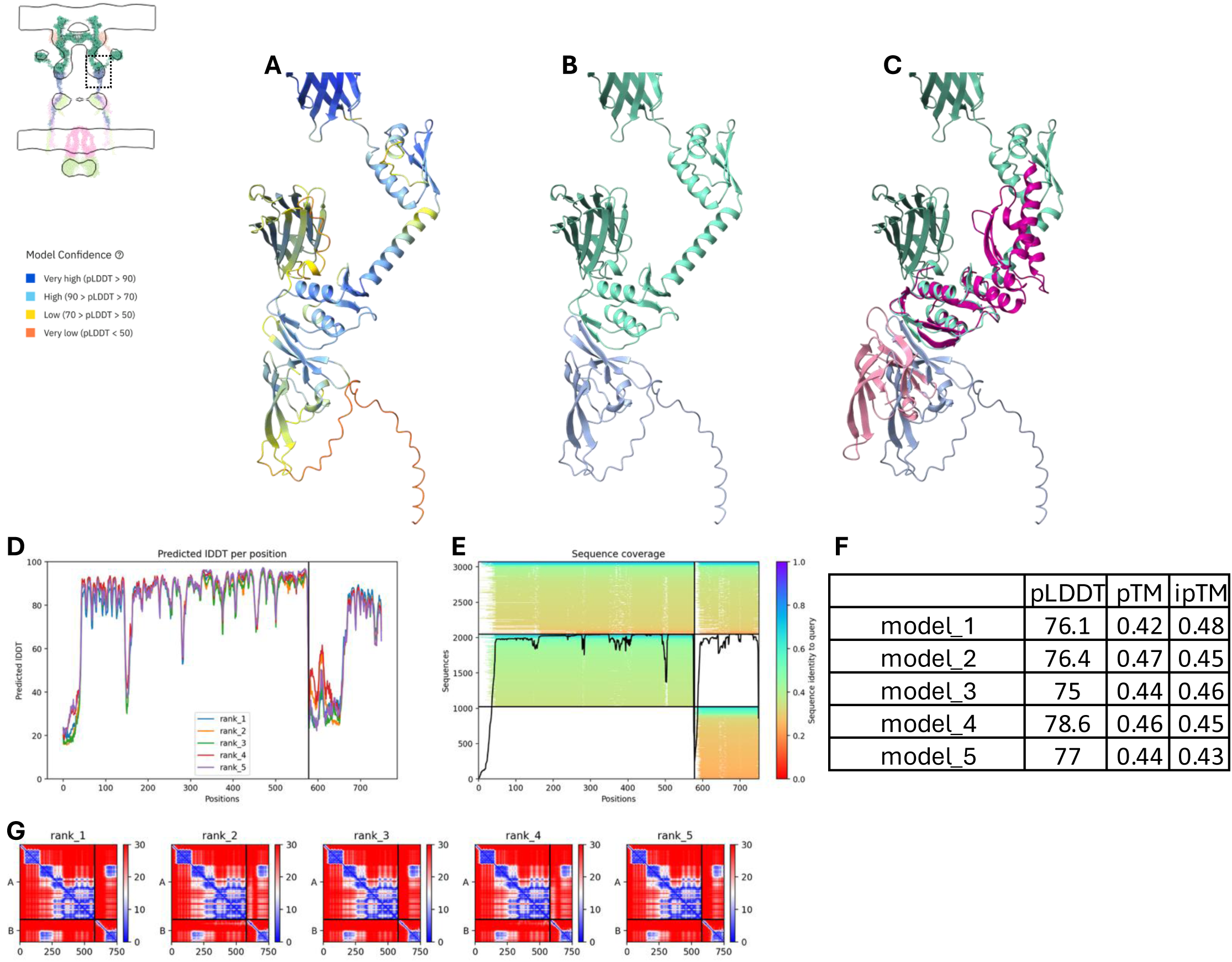
PilQ-PilP model. A) AF prediction of the interaction between PilP and the PilQ Secretin/TonB short N-terminal domain colored by pLDDT or B) by chain identity. C) Superimposition of the PilQ-PilP model with the *Neisseria meningitidis* solved PilQ-PilP complex (4AV2). D) AF predicted local Distance Difference Test, E) Sequence coverage, F) summary table of the 5 models produced by ColabFold G) Predicted Aligned Error matrix (colormap in Ångströms).

**Figure S9.**
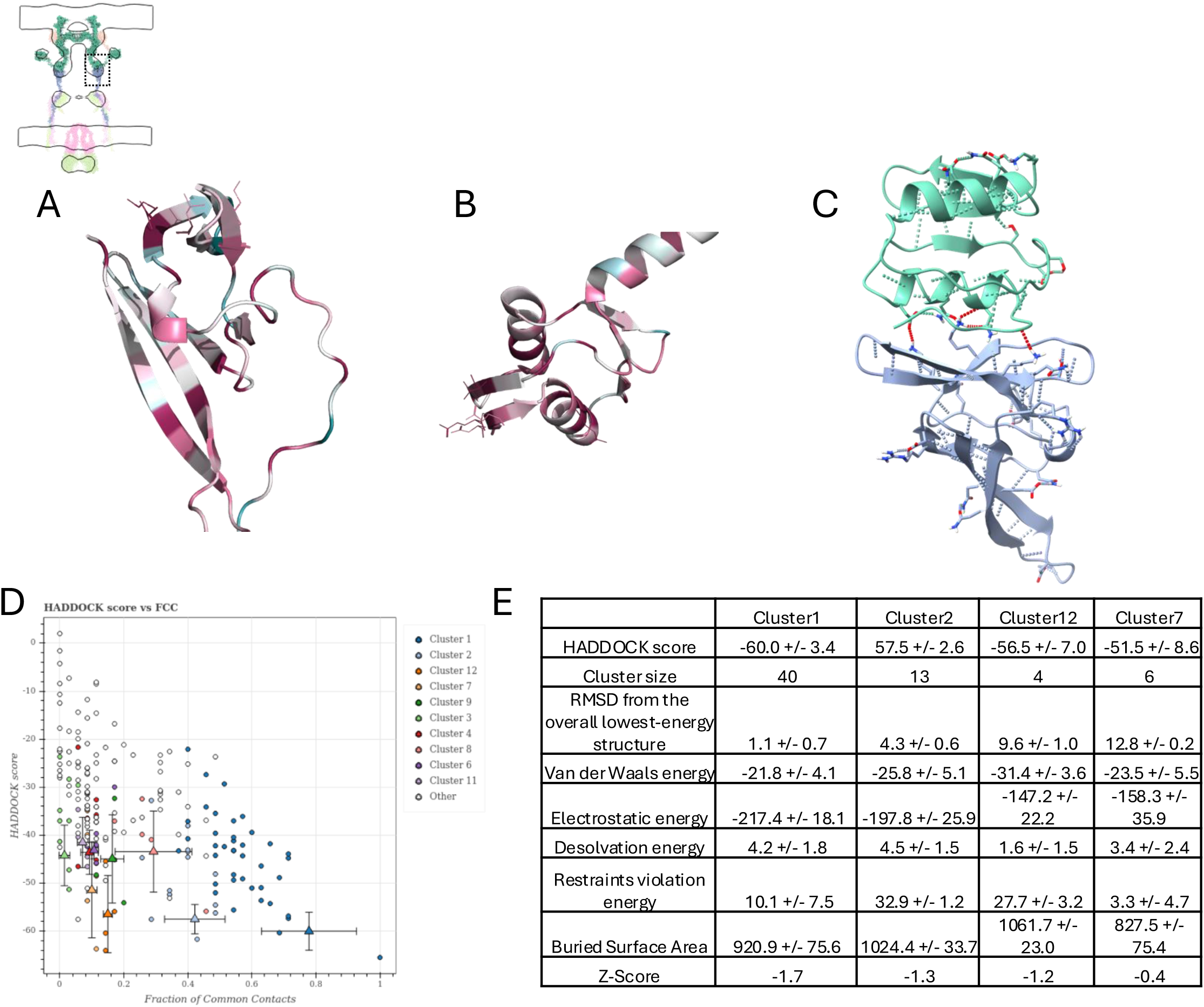
Conservation-driven HADDOCK model for PilQ-PilP interaction. Based on the *Nm*PilQ-PilP structure (4AV2), we identified conserved residues (using Consurf) in the interacting interface. A) PilP: L117, K105, K127. B) PilQ: N162, Q164, D165. C) HADDOCK docking, D) cluster scores, and E) statistics table using the previously identified residues.

**Figure S10.**
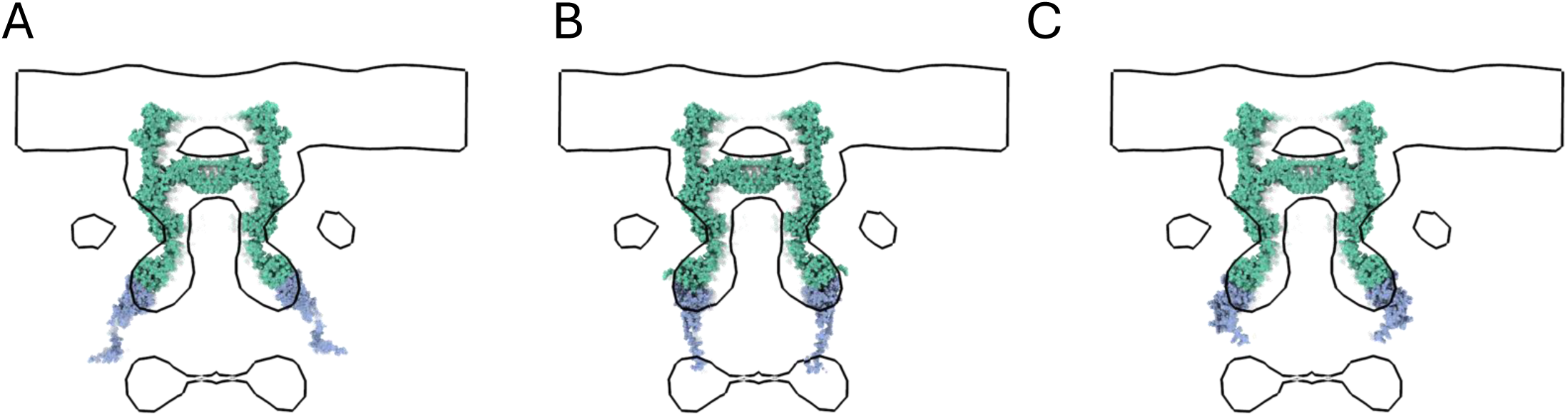
Fitting of different models into the non-piliated STA. We tested three different models for the PilQ-PilP interaction using A) the *Nm*PilQ-PilP structure (4AV2) to dock PilP on PilQ, B) the conserved residues at the Protein-Protein interface to dock PilP on PilQ using HADDOCK, and C) the ColabFold PilQ-PilP model. Model B seems to better occupy the secretin density and the connectivity with the lower periplasmic ring.

**Figure S11.**
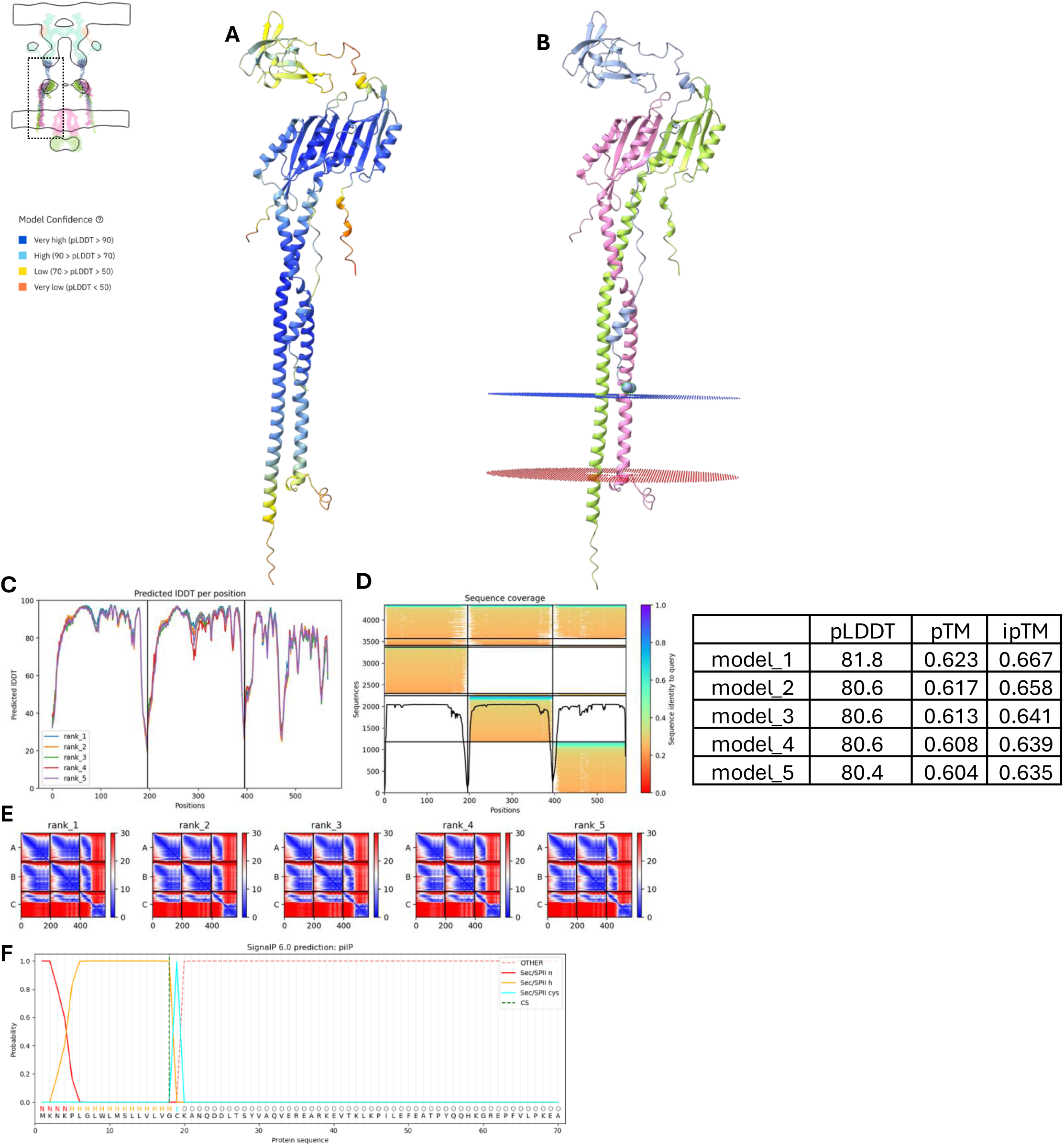
PilN-O-P ColabFold model and membrane insertion. A) AF prediction of the PilN-O-P interaction. B) This model is well-positioned to insert into the membrane, and the predicted lipidated cysteine (showed as a ball) on PilP is in the right position. C) AF predicted local Distance Difference Test, D) Sequence coverage, and E) Predicted Aligned Error matrix (colormap in Ångströms). F) SignalP prediction of cysteine 19 lipidation.

**Figure S12.**
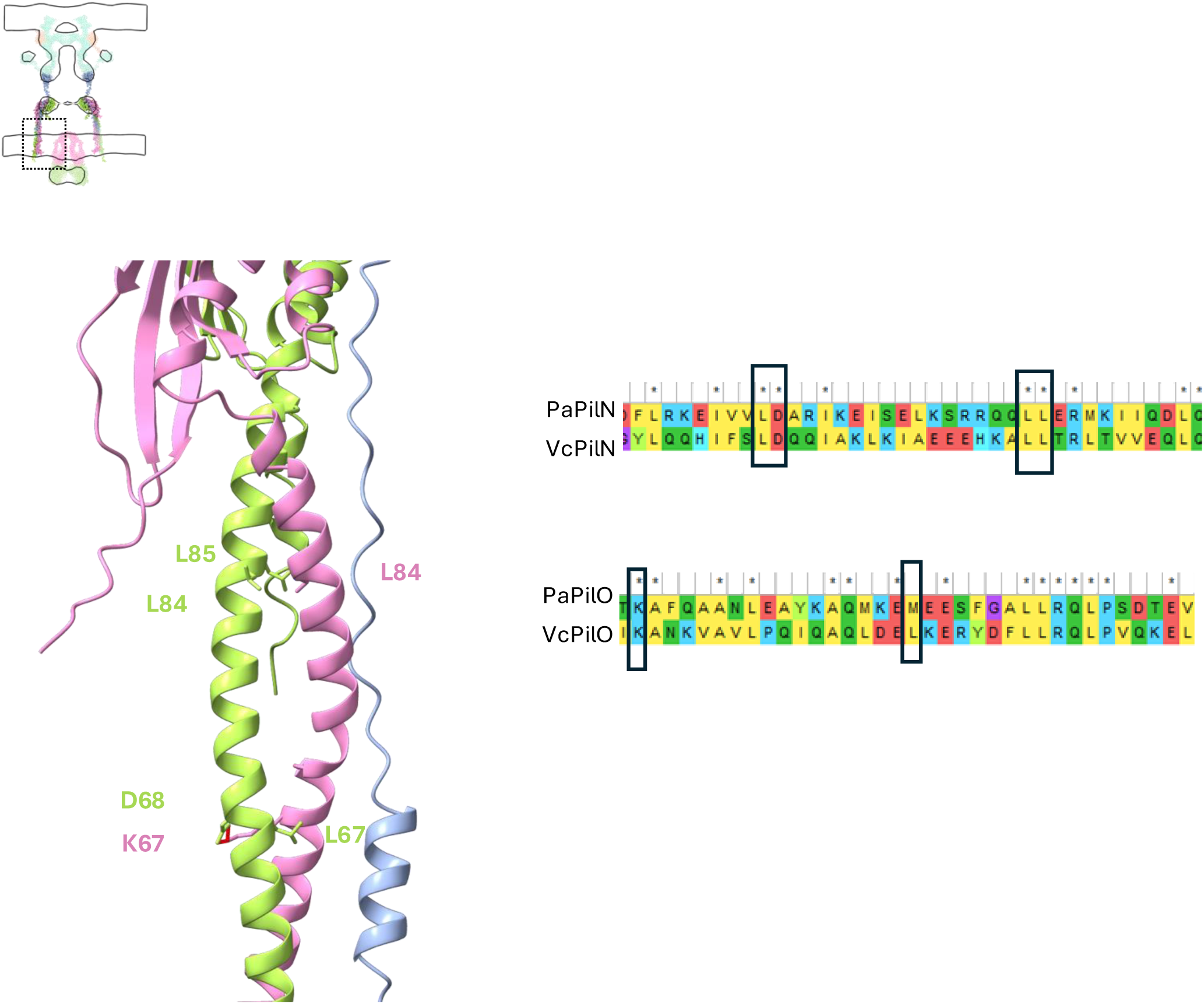
ColabFold PilN-O-P agrees with mutagenesis data. Locations of residues involved in PilN-O interaction identified in the *Pseudomonas aeruginosa* homologs^31^. AlphaFold predicted a salt bridge between PilO K67 and PilN D68, and these residues are conserved in *Pseudomonas aeruginosa*.

**Figure S13.**
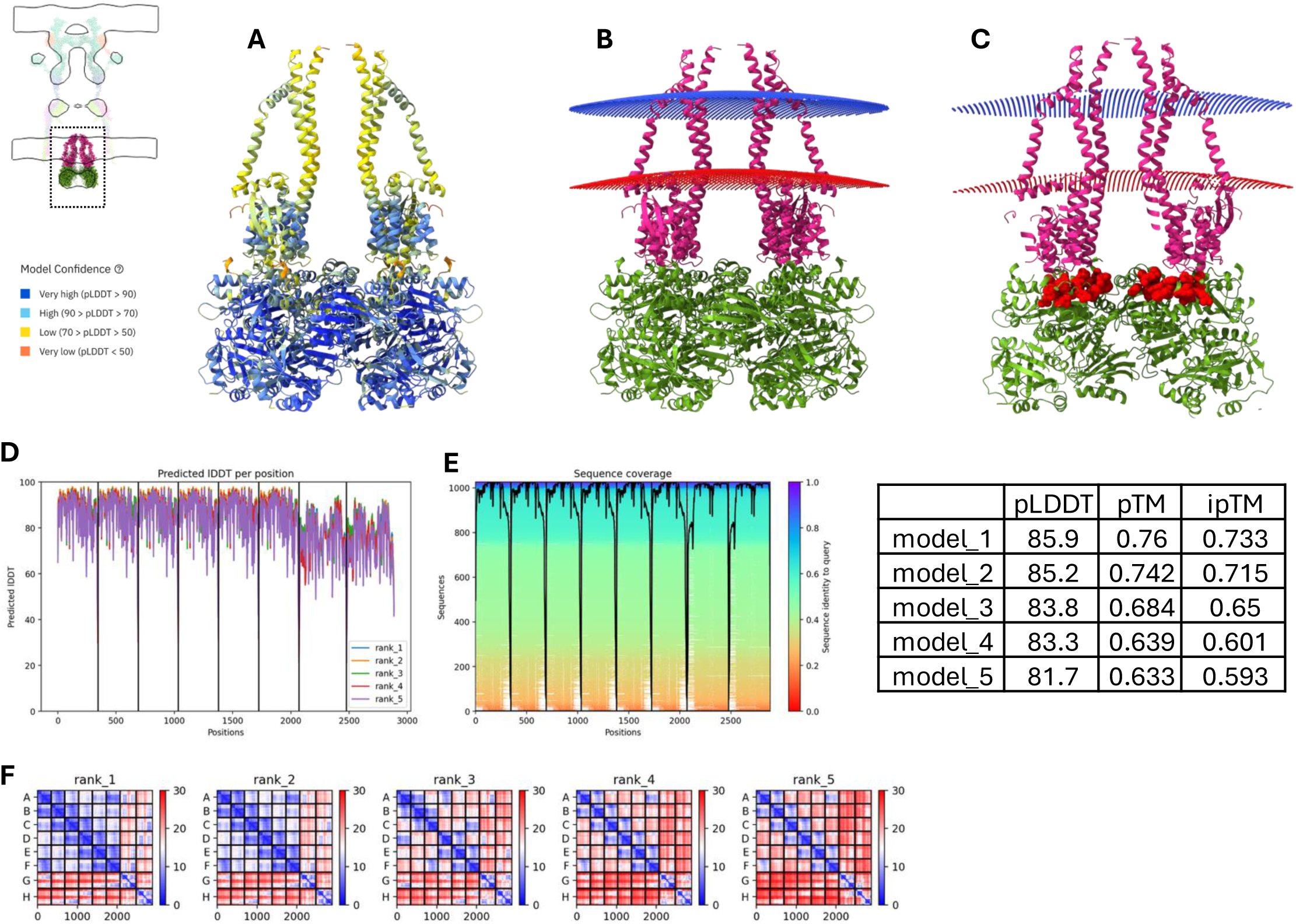
ColabFold PilC-PilT model agrees with mutagenesis data. A) AF prediction of the PilC-PilT interaction. B) This model is well-positioned to insert into the membrane. C) The location of the PilT AIRNLIRE motif (rendered as red spheres) at the PilC interface could explain why this motif is required for retraction but not for ATPase activity. D) AF predicted local Distance Difference Test, E) Sequence coverage, and F) Predicted Aligned Error matrix (in Ångströms).

**Figure S14.**
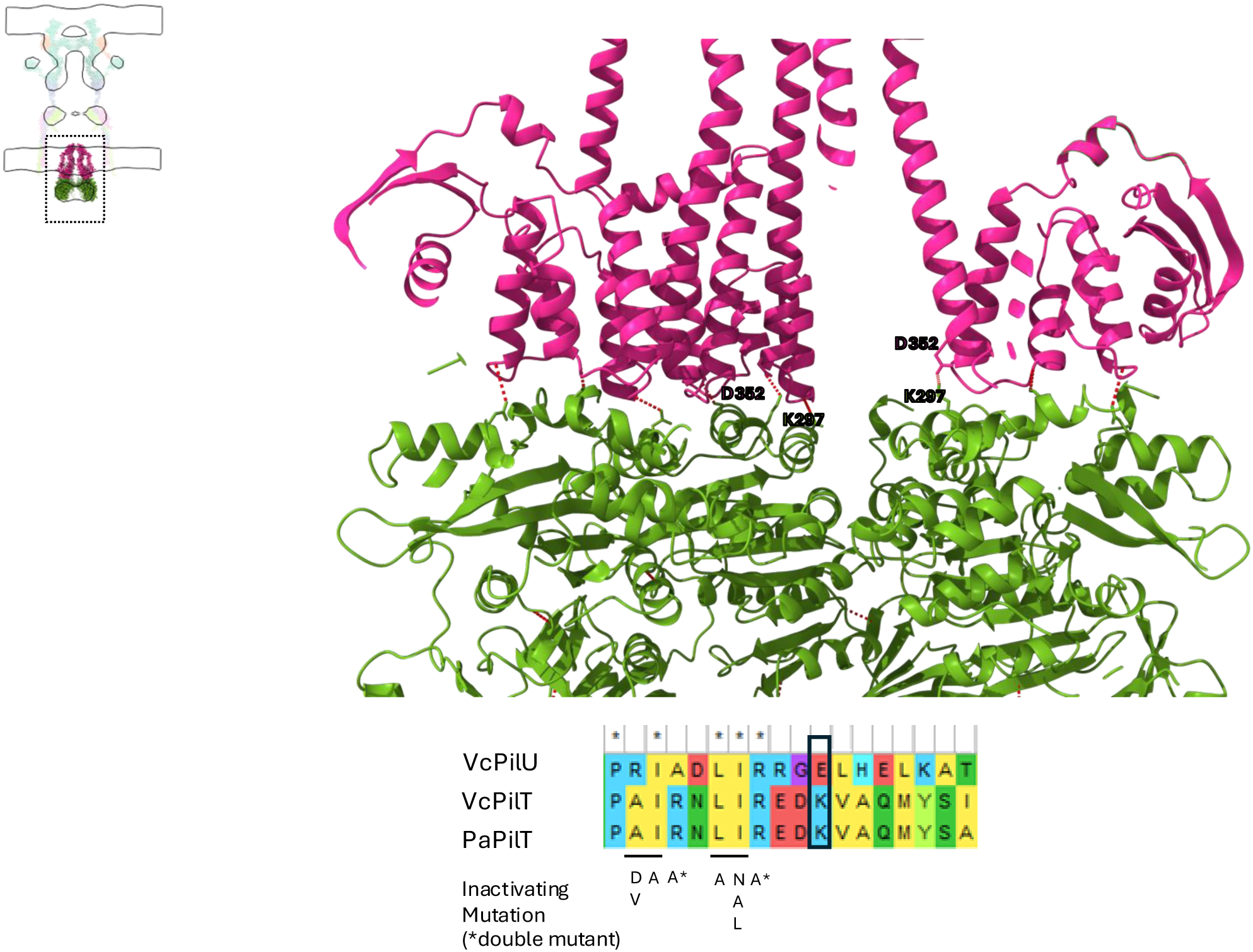
ColabFold PilC-PilT model agrees with mutagenesis data. Top) Hydrogen bond network between PilC and PilT models with a focus on PilC D352 and PilT K297. Bottom) MSA of PilU, PilT, and the biochemically characterized *Pa*PilT showing residues important for pili formation but not involved in ATP hydrolysis.

**Figure S15.**
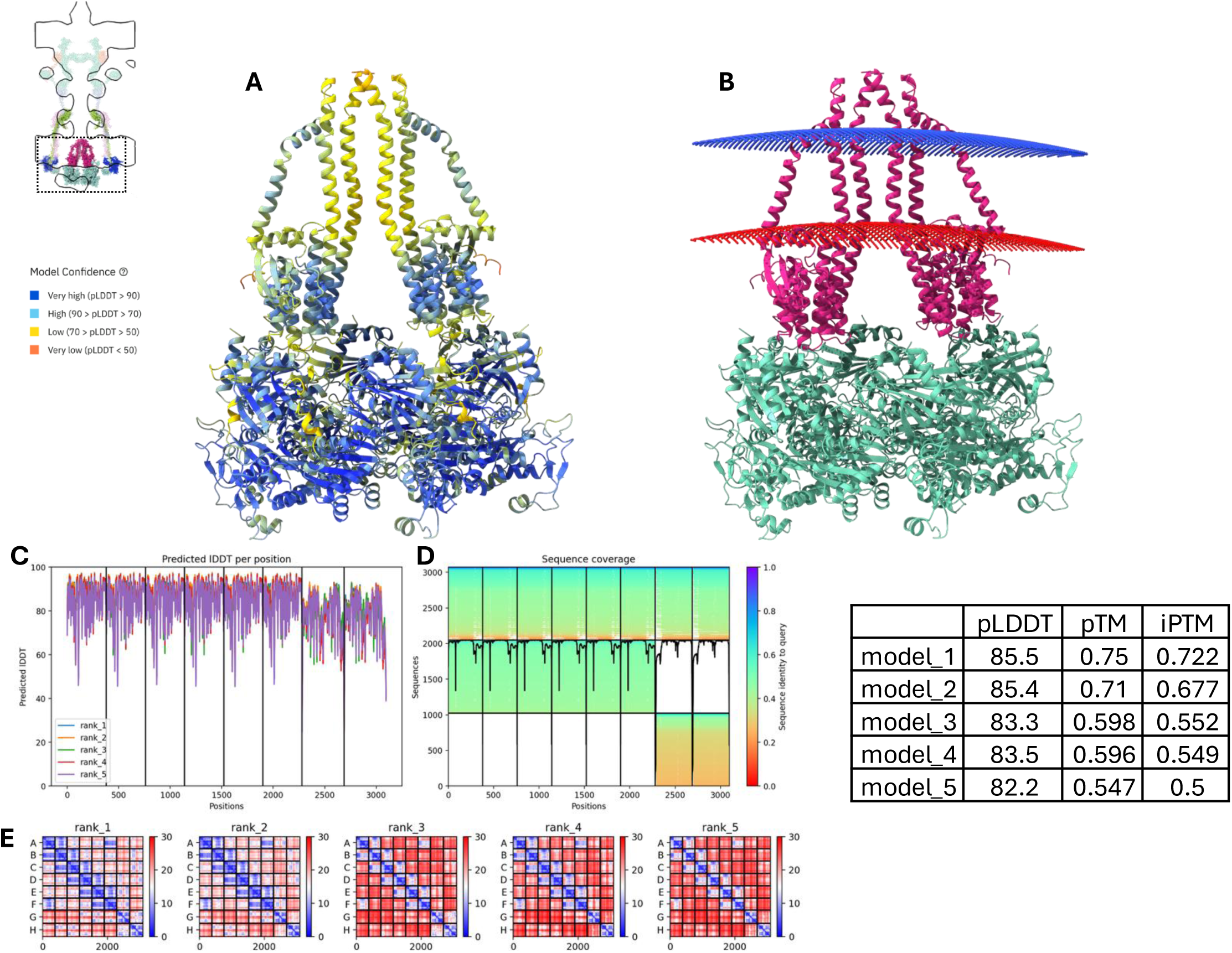
ColabFold PilB-C model. A) AF prediction of the PilB-PilC interaction. B) This model is well-positioned to insert into the membrane. C) AF predicted local Distance Difference Test, D) Sequence coverage, and E) Predicted Aligned Error matrix (in Ångströms).

**Figure S16.**
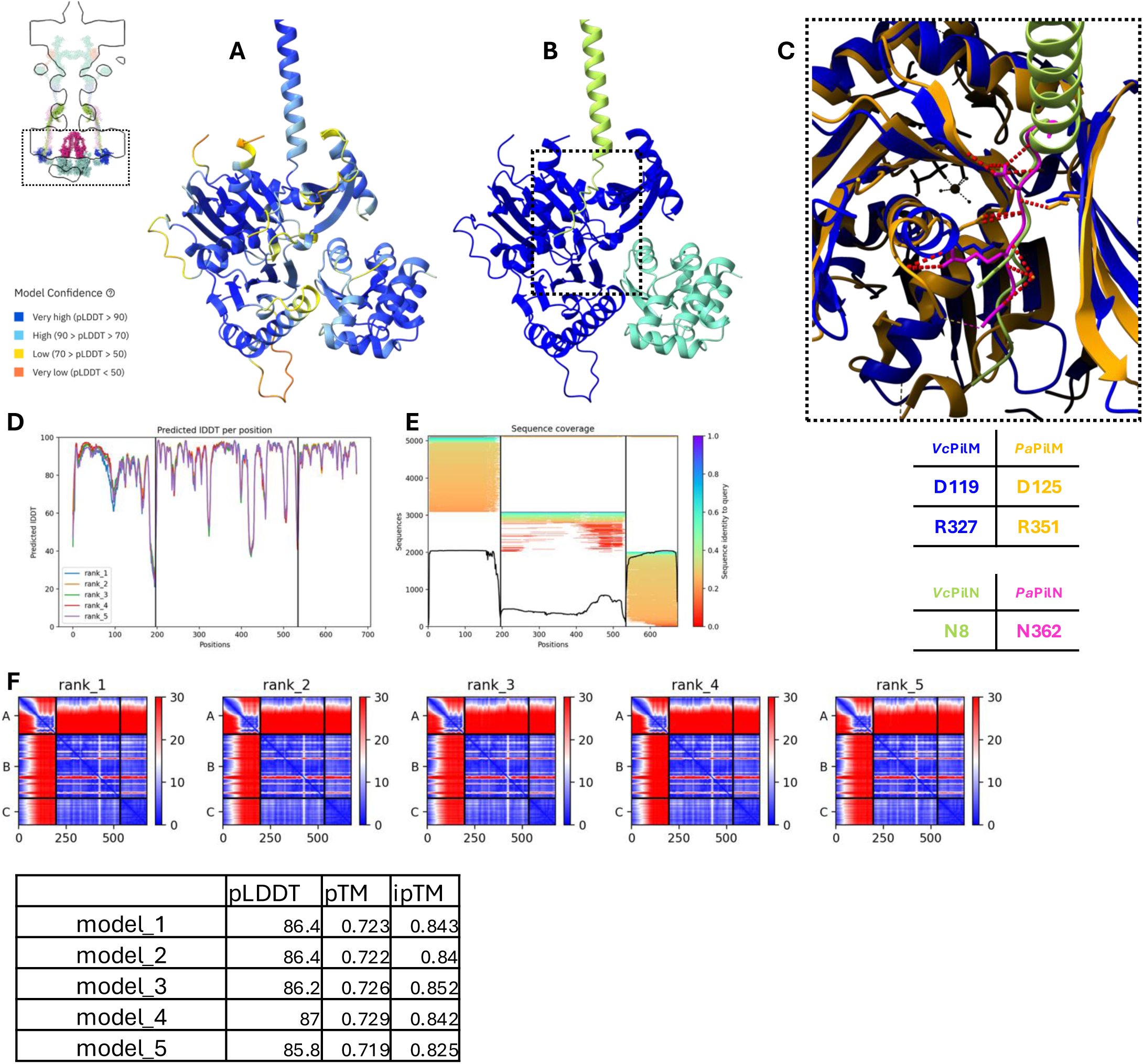
ColabFold can predict PiN-PilM and PilM-PilB interactions. A) AF prediction of the PilN-PilM-PilB interactions. B) Model in (A) colored by chain. C) Enlarged views of the PilN-PilM binding region. The AF model closely resembles the *Pa*PilM-N complex’s solved structure (PDB:5EOU) at a residue level. D) AF predicted local Distance Difference Test, E) Sequence coverage, and F) Predicted Aligned Error matrix (in Ångströms).

**Figure S17.**
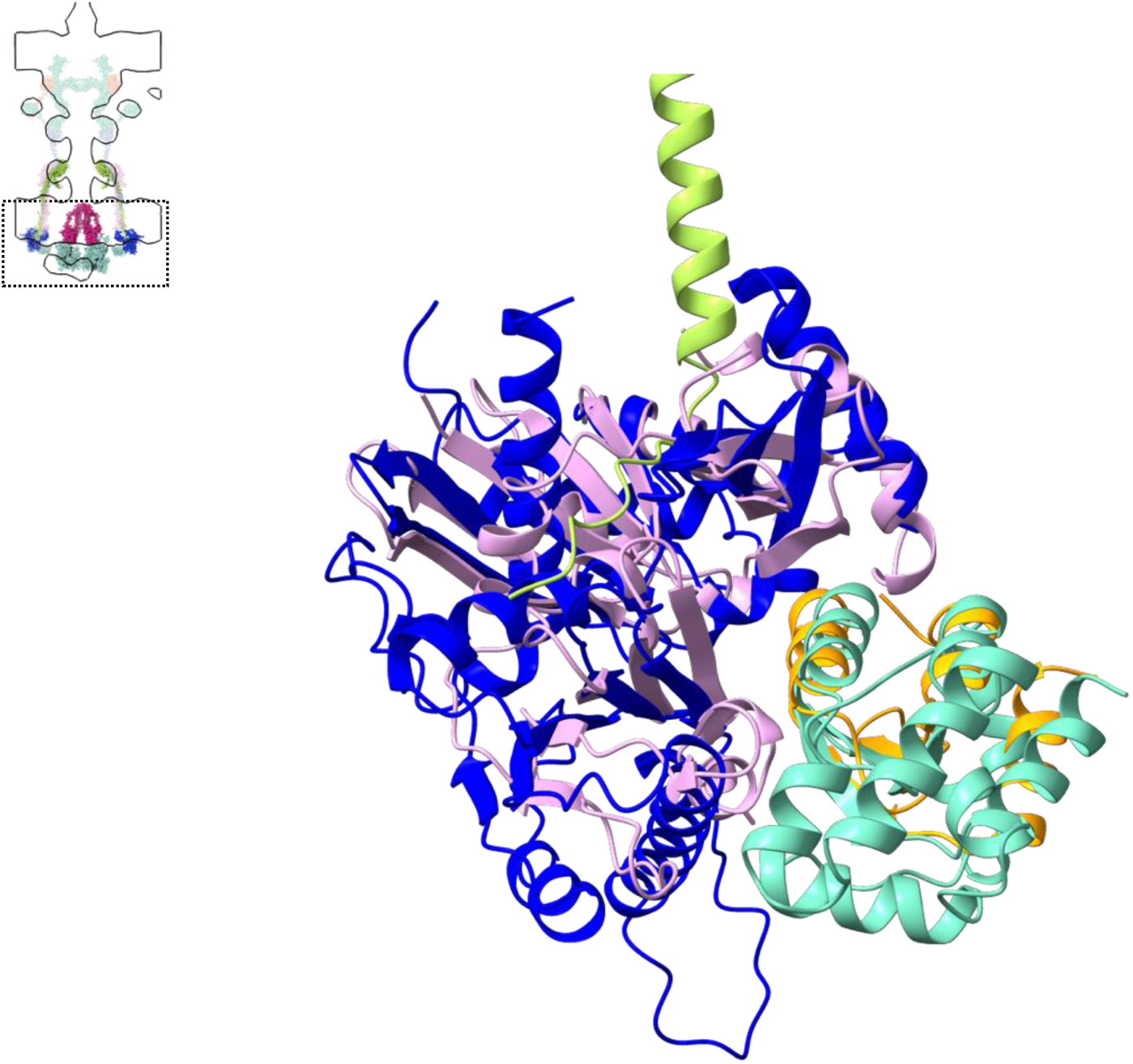
ColabFold can predict PilB’s binding region on PilM. A) Superimposition of the PilN-PilM-PilB AF multimer model (colored in lime green, blue and turquoise) with the solved *Vc*EspL-EspE (colored in pink and orange respectively) structure (PDB: 2BH1) from the closely related type II secretion system (sequence alignment score = 158.2; RMSD between 68 pruned atom pairs is 1.3 Å; across all 211 pairs: 15.2 Å). AF appears to correctly predict the binding interface.

**Figure S18.**
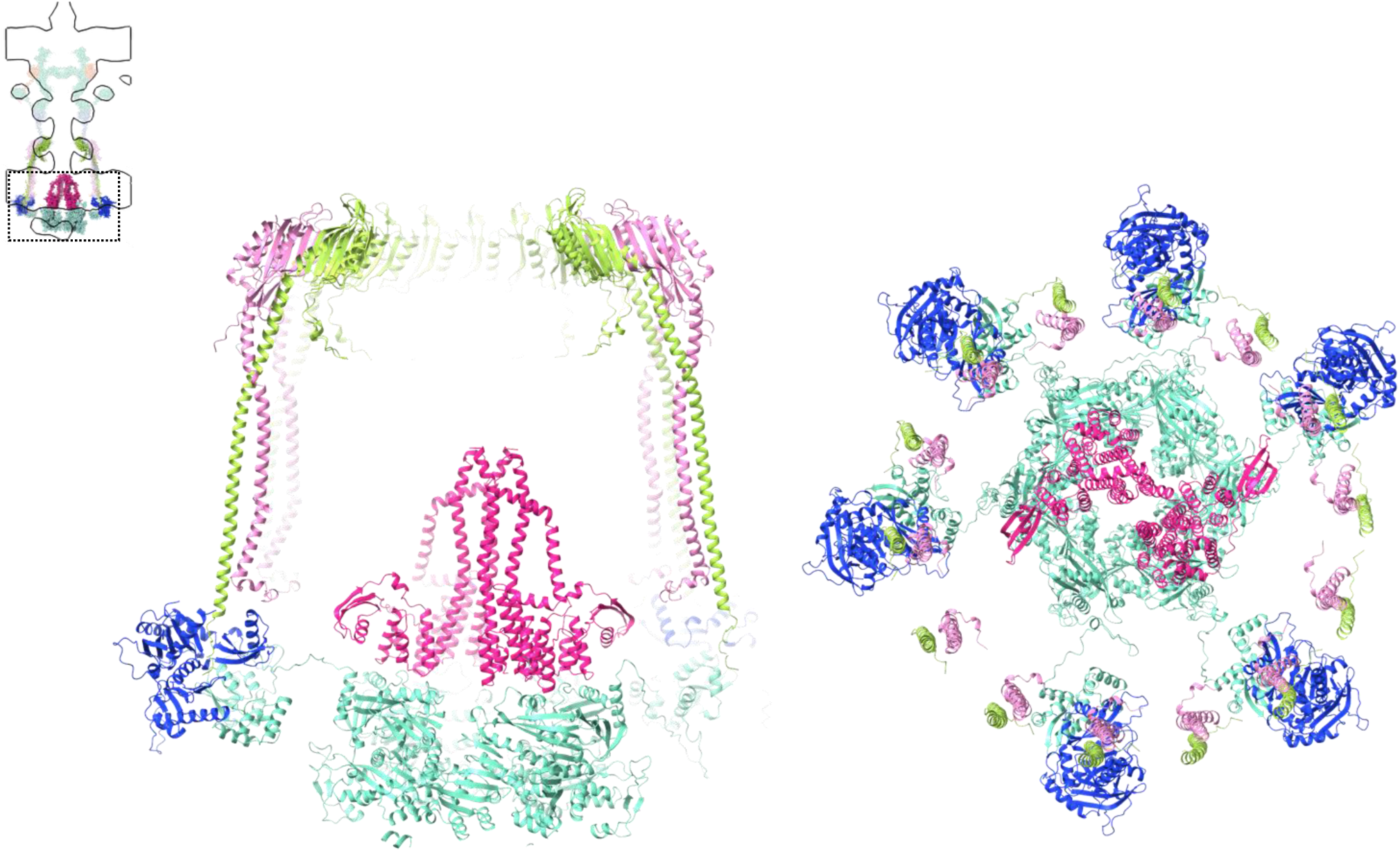
Complete model of the inner membrane platform. Final model of the inner membrane platform obtained by combining the PilN-PilO-PilP, PilM-PilN-PilB, and PilB-PilC models.

**Figure S19.**
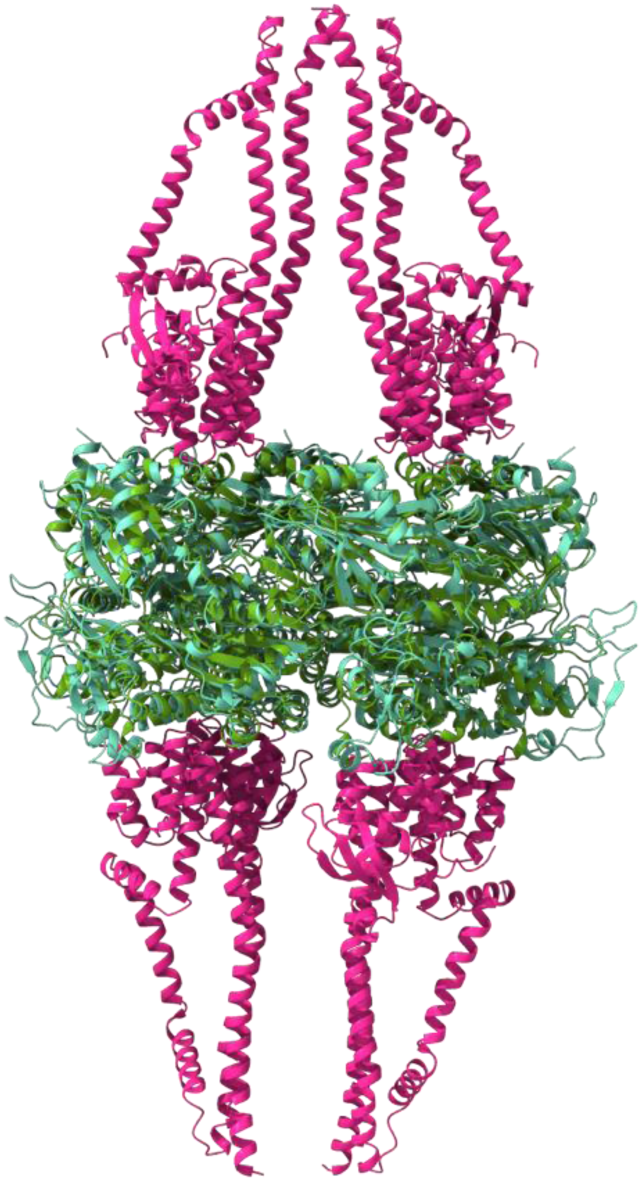
PilB and PilT could bind PilC rotated 180°. Superimposition of the PilB-C and the PilT-C models reveal PilB and PilT likely interact with PilC using two different interfaces (sequence alignment score = 576.5; RMSD between 228 pruned atom pairs is 0.980 Å; across all 311 pairs: 2.906 Å).

**Figure S20.**
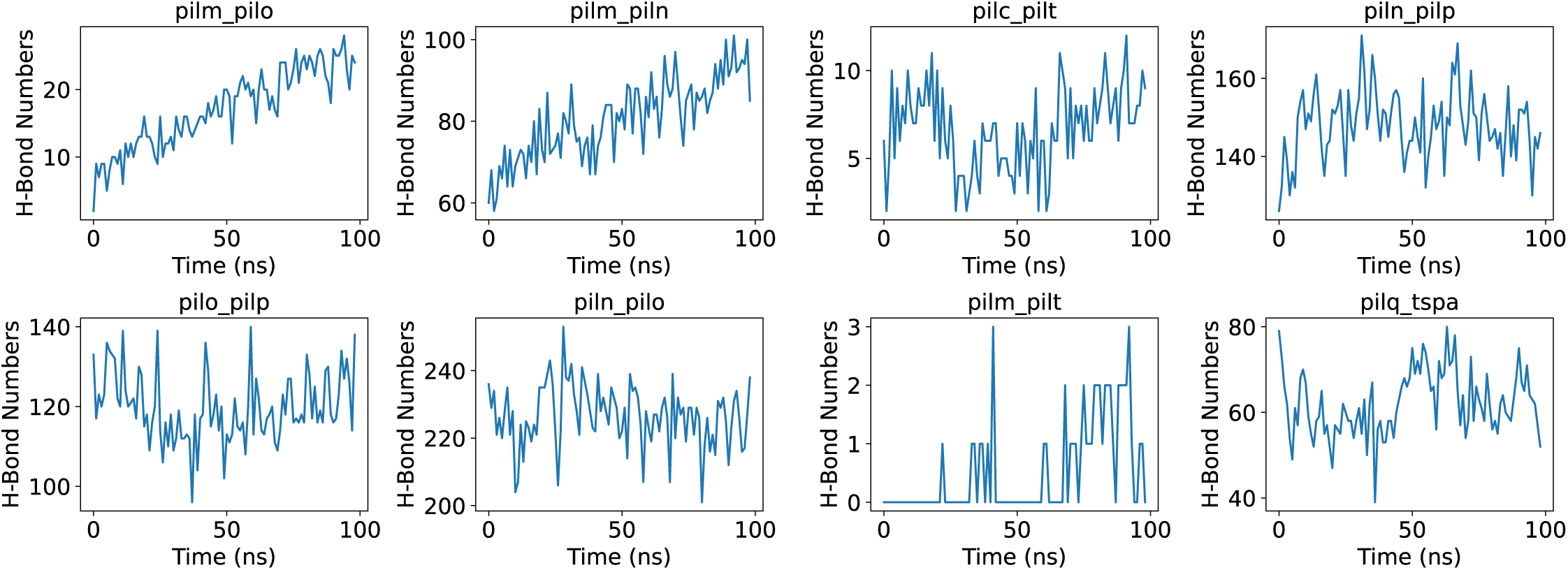
H-Bond formation between different pairs of components during MD equilibration.

**Figure S21.**
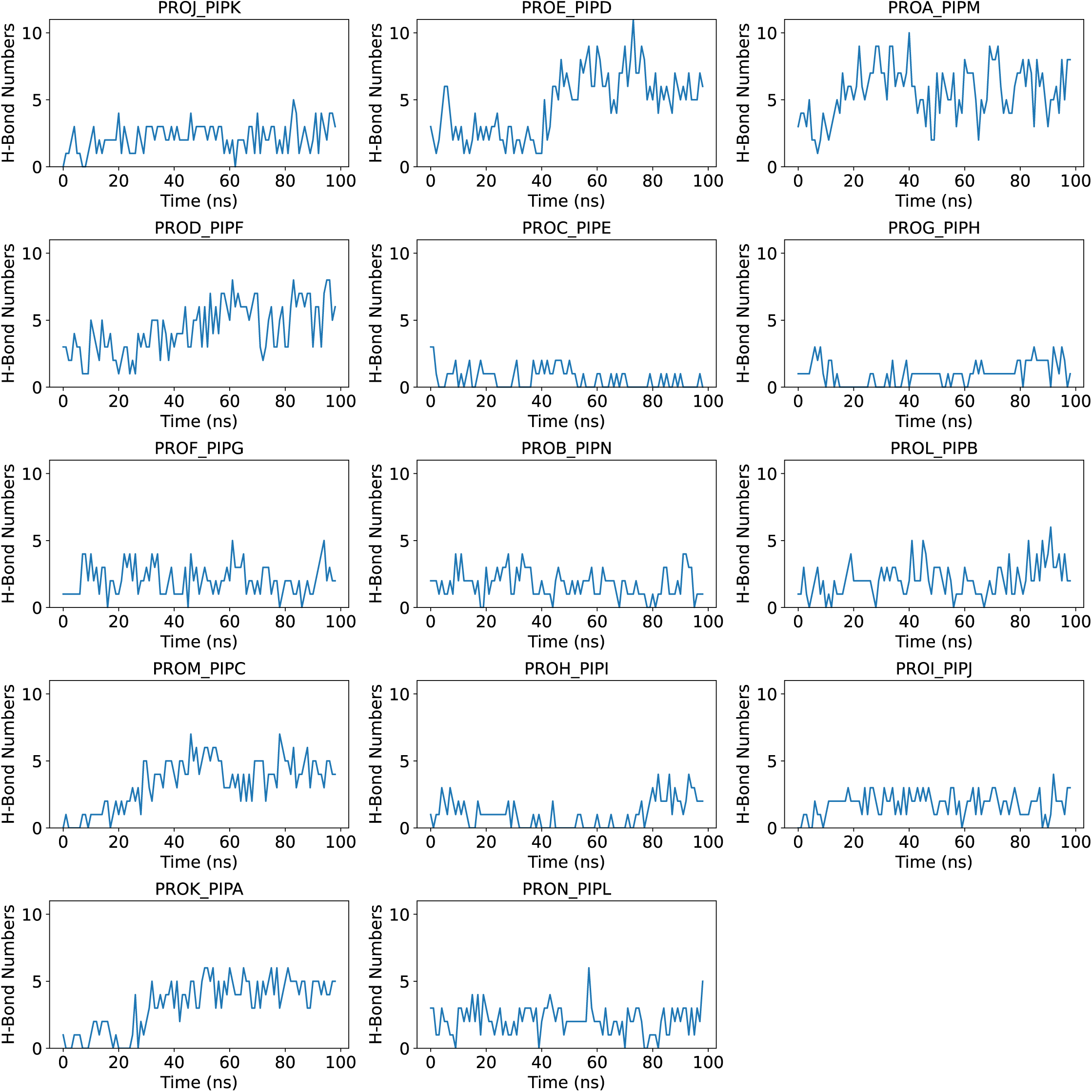
Number of H-Bonds between different individual pairs of PilP and PilQ during MD equilibration.

## Strains used in this paper

| Strain # | Genotype | Application |
| --- | --- | --- |
| SAD2096 | E7946 Sm<sup>R</sup>, P<sub>tac</sub>-tfoX, P<sub>tac</sub>-recA, ΔlacZ::lacIq, comEA-mCherry, ΔCTX::Kan<sup>R</sup>, ΔMSHA::Carb<sup>R</sup>, ΔluxO, ΔTCP::Zeo<sup>R</sup>, ΔpilT::Tm<sup>R</sup> | pili reconstruction and CryoET |
| TND3651 / SAD3999 | E7946 Sm<sup>R</sup>, P<sub>tac</sub>-tfoX, P<sub>tac</sub>-recA, ΔlacZ::lacIq, comEA-mCherry, ΔCTX::Kan<sup>R</sup>, ΔMSHA::Carb<sup>R</sup>, ΔluxO, ΔTCP::Zeo<sup>R</sup>, msfGFP inserted into pilQ after T35 | PilQ AMIN domain localisation |
| TND3601 / SAD4000 | E7946 Sm<sup>R</sup>, P<sub>tac</sub>-tfoX, P<sub>tac</sub>-recA, ΔlacZ::lacIq, comEA-mCherry, ΔCTX::Kan<sup>R</sup>, ΔMSHA::Carb<sup>R</sup>, ΔluxO, ΔTCP::Zeo<sup>R</sup>, pilT<sup>K136A</sup> ΔpilU::Tm<sup>R</sup> | PilT localisation |
| TND3780 / SAD4001 | E7946 Sm<sup>R</sup>, P<sub>tac</sub>-tfoX, P<sub>tac</sub>-recA, ΔlacZ::lacIq, comEA-mCherry, ΔCTX::Kan<sup>R</sup>, ΔMSHA::Carb<sup>R</sup>, ΔluxO, ΔTCP::Zeo<sup>R</sup>, pilT<sup>K136A</sup> pilU<sup>K134A</sup> | PilU localisation |
| TND0905 / SAD2468 | E7946 Sm<sup>R</sup>, ΔlacZ::lacIq, P<sub>tac</sub>-tfoX, ΔluxO, pilA<sup>S67C</sup> | Transformation, fluorescence microscopy, and aggregation assays |
| TND3660 / SAD4002 | E7946 Sm<sup>R</sup>, ΔlacZ::lacIq, P<sub>tac</sub>-tfoX, ΔluxO, ΔpilA::Kan<sup>R</sup> | Transformation, fluorescence microscopy, and aggregation assays |
| AET133 / SAD3996 | E7946 Sm<sup>R</sup>, ΔlacZ::lacIq, P<sub>tac</sub>-tfoX, ΔluxO, pilA<sup>S67C,K40S</sup> | Transformation, fluorescence microscopy, and aggregation assays |
| AET134 / SAD3997 | E7946 Sm<sup>R</sup>, ΔlacZ::lacIq, P<sub>tac</sub>-tfoX, ΔluxO, pilA<sup>S67C,K108S</sup> | Transformation, fluorescence microscopy, and aggregation assays |
| AET135 / SAD3998 | E7946 Sm<sup>R</sup>, ΔlacZ::lacIq, P<sub>tac</sub>-tfoX, ΔluxO, pilA<sup>S67C,K120S</sup> | Transformation, fluorescence microscopy, and aggregation assays |

