## Supplemental Modeling Information for "Integrative Structure and Function of the *Vibrio cholerae* Competence Pilus Machine"

To build the CPM model, we used a combination of AI-generated models and solved structures as templates to model protein-protein interactions while using the STA map of the $PilT_{K136A}$ $PilU_{K133A}$ strain as an envelope. Recent benchmarking of deep learning methods to predict structures of protein complexes showed that AlphaFold and ColabFold[1] perform on a similar level and that a confidence score (ipTM) of 0.75 could be used as threshold for discerning good from poor complexes[2].

Of all the CPM components, only the *Vc*PilQ structure has been solved, but homologous protein complexes have been solved in other species, such as *Pseudomonas aeruginosa* and *Nesseria meningitis*. To model the *Vc*PilQ-TsaP interactions, we used the solved *Pa*PilQ-TsaP (PDB: 6VE2) as a starting point. Superimposition of *Vc*PilQ to *Pa*PilQ led to a good alignment (RMSD=1.1 Å between 142 pruned atom pairs and 7.2 Å across all 294 pairs), so we then used *Pa*TsaP as template to dock the *Vc*TsaP AlphaFold model (Fig. S8-9, RMSD=1.3 Å between 71 pruned atom pairs and 5.7 Å cross all 206 pairs). We then refined *Vc*TsaP with C7 symmetry consistent with the *Pa*TsaP solved structure.

To model the PilQ-PilP interaction, we tested several strategies: first, we submitted PilQ and PilP to ColabFold (Fig. S10). The predicted ColabFold models have a maximum ipTM score of 0.481, where PilQ and PilP interact in a way resembling the crystal structure of the type II secretion system's GspC-GspD complex (PDB:3OSS) (Fig. S10). Second, we used the cryo-EM structure of the *Nm*PilQ-PilP complex (PDB: 4AV2) as a template to guide the docking (Fig. S11). The model resulting from using the *Nm*PilQ-PilP complex as an envelope shows that PilP has a RMSD of 1.180 Å (for 47 atoms; for all 81 pairs, RMSD=2.6 Å), but when fitted into our STA, PilP seems to be oriented in a wrong orientation based on its position relative to the lower periplasmic ring (Fig. S12). We used HADDOCK to refine PilP's position using conserved residues as docking points (Fig. S11). After the docking, a series of hydrogen bonds are formed at the protein's interface (Fig. S11C). This last model fits to our map better, with PilP residues pointing to the lower periplasmic ring in a more relaxed way (Fig. S12).

We then manually fitted the PilQ-TsaP-PilP complex into the non-piliated map. We retrieved the full-length PilQ model from UNIPROT, docked its AMIN domain into the secretin's flanking density, and rebuilt the connecting loop using Modeller[3]. Finally, we symmetrized the full-length PilQ model to match the solved structure.

For the $PilN_1PilO_1PilP_1$, $PilN_1PilM_1PilB_1$, $PilC_2PilB_6$, $PilC_2PilT_6$, and $PilU_6$ complexes, we used ColabFold multimer with standard parameters. Because there is no solved structure of the $PilN_1PilO_1PilP_1$ assembly, we tested the topology of the assembly by in-silico membrane insertion. After membrane insertion, the predicted PilN-PilO transmembrane segments and the predicted lipidated PilP cysteine are correctly positioned (Fig. S13). We further tested the quality of the model using the extensively characterized *Pa*PilN-PilO interaction[4]. Residues important for *Pa*PilN-PilO coil-coil interactions (*Pa*PilN: L82, L81, L64, D65; *Pa*PilO: L85, M92, L67, D68; and *Vc*PilO: L85, L84, L67, D68) are conserved in *V. cholerae* and, in our model, are localized at the protein-protein interface. In particular, *Vc*$PilN^{L85}$, *Vc*$PilN^{L84}$, and *Vc*$PilO^{L84}$ are arranged in leucine zipper motif (Fig. S14).

We checked the quality of our $PilN_1PilM_1PilB_1$ model by comparison with the *Pa*$PilN_{1-12}$-PilM complex[11] (PDB: 5EOU, Fig. S15). The two models superimpose well (5.2 Å RMSD across all 323 atoms pairs) and have a similar network of interacting residues between PilN and PilM. To confirm the PilM-PilB interactions, we superimposed our model to the EspL-EspE complex from the homologous type II secretion system[12] in *E.*

*coli* (PDB: 2BH1, Fig. S16). Even though *Vc*PilM and *Ec*EspL don't share high sequence similarity, in our model *Vc*PilB and *Ec*EspE have an overall similar orientation (RMSD between 68 pruned atom pairs is 1.272 Å; across all 211 pairs: 15.226 Å). These structure details, combined with a high ipTM score, suggest that the predicted model could be correct.

Multiple structures are available for the three ATPases of *V. cholerae* agree on a hexameric nature of PilB, PilT, and PilU, but the ATPase-PilC complex is not solved yet[5,6]. In our $PilC_2PilT_6$ model, two salt bridges were identified (linking $PilT_{K297}$ with $PilC_{E352}$ and $PilT_{K297}$ with $PilC_{D150}$). In *Pa*PilT, a mutation of K297 abolishes twitching mobility, highlighting the importance of this salt bridge in type IV pilus dynamics[7]. Moreover, the AIRNLIRE motif, which is required for retraction but not for ATPase activity, is nearby the interacting interface (Fig. S18–19). This, combined with a good ipTM score, highlights the high quality of the model.

In our $PilC_2PilB_6$ model (Fig. S20), PilC interacts with PilB through its PAS-like N-terminal domain, consistent with evidence that this domain rotates during transition from PilC's open to closed conformation in *Thermus thermophilus*[8] and in line with the *Xanthomonas citri* type IV inner membrane platform model[9]. Unlike PilC-PilT, no specific contact points were identified, but the interaction is in line with previous evidence showing electrostatic interactions between PilB and PilC[10]. Lastly, we built the loop connecting PilB's ATPase domain with PilM using Modeller.
